# Unsupervised discovery of dynamic cell phenotypic states from transmitted light movies

**DOI:** 10.1101/2021.01.26.428210

**Authors:** Phuc H.B. Nguyen, Sylvia Chien, Jin Dai, Raymond J. Monnat, Pamela S Becker, Hao Yuan Kueh

## Abstract

Identification of cell phenotypic states within heterogeneous populations, along with elucidation of their switching dynamics, is a central challenge in modern biology. Conventional single-cell analysis methods typically provide only indirect, static phenotypic readouts. Transmitted light images, on the other hand, provide direct morphological readouts and can be acquired over time to provide a rich data source for dynamic cell phenotypic state identification. Here, we describe an end-to-end deep learning platform, UPSIDE (for Unsupervised Phenotypic State IDEntification), for discovering cell states and their dynamics from transmitted light movies.

UPSIDE uses the variational auto-encoder architecture to learn latent cell representations, that are then clustered for state identification, decoded for feature interpretation, and linked across movie frames for transition rate inference. Using UPSIDE, we identified distinct blood cell types in a heterogeneous dataset. From acute myeloid leukemia cell movies, we then identified stem-cell associated morphological states and their inter-conversion rates. UPSIDE opens up use of transmitted light movies for systematic exploration of cell state heterogeneity and dynamics in biology and medicine.

## Introduction

Cells maintain and switch between distinct phenotypic states in a dynamic manner. The facile identification of these states, and understanding the basis for and dynamics by which they interconvert, is a central challenge in biology. Modern single-cell analysis methods, such as single cell RNA sequencing and multiparameter flow cytometry or mass cytometry^1–5^, are widely used to define cell states in heterogeneous populations; while powerful, these methods provide incomplete readouts of cell phenotypes, and typically do not report on stability or transition dynamics. Transmitted light microscopy images directly reveal cell morphology, and have historically formed the basis for identifying cell types and cell states in diverse fields, ranging from cell biology to neuroscience^6, 7^. These images can then be acquired at successive timelapse intervals and over long times, with minimal phototoxicity and without prior labeling or genetic manipulation. The resultant live cell movies can reveal additional information about the dynamics of these cell phenotypic states.

Cell phenotypes have traditionally been identified by the visual inspection and interpretation of transmitted light or electron micrographic images. The advent of modern machine learning, however, is enabling high-throughput automated analysis of cell morphology, and could open doors for new deep learning approaches for the systematic, unbiased extraction of cell morphological states and their transition dynamics from these imaging data sets. However, several barriers remain to the development of such methods. First, current deep learning pipelines for cell image analysis rely heavily on predetermined knowledge to generate classification training datasets, or on large sets of heuristic formulations to capture the diversity of cell shapes and morphologies ^8–11^. When examining novel biological processes with minimal to no preconceived information, it can be difficult for investigators to determine what the important labels are without manual intervention and feature selection. Second, current machine learning pipelines generate features that are often not readily interpretable. A variety of unsupervised methods can generate reduced dimensionality representations from complex data, including principal component analysis (PCA), adversarial autoencoders^12^, generative adversarial network ^13, 14^, and self-supervised deep learning approaches ^15, 16^. However, these methods are limited in their ability to generate interpretable morphological features that allow scientists to investigate and understand the machine-identified cell states. Finally, current movie analysis methods cannot infer state transition dynamics from live cell movies in an automated, systematic manner^17^. Cell state transitions are typically observed from trajectories of single cells; however, despite recent advances^18^, current tracking algorithms still typically require considerable parameter adjustment and manual error correction for generation of cell trajectories^19^.

Here, we present an end-to-end deep learning method for elucidating cell phenotypic states and their dynamics from brightfield movies of living cells. This method, termed UPSIDE (for Unsupervised Phenotypic State IDEntification), is designed to facilitate unsupervised discovery of cellular phenotypic states, elucidation of morphological features that define these states, and inference of state transition dynamics. UPSIDE segments cells directly from brightfield images, then utilizes the variational autoencoder architecture (VAE)^20^ to learn intuitive latent features that can be clustered to reveal distinct morphological states, and also decoded to extract human-interpretable meaning. In order to demonstrate use and versatility of UPSIDE, we first used distinct hematopoietic cell phenotypes in a mixed dataset. We then analyzed live imaging movies of leukemic cells from an acute myeloid leukemia (AML) patient to identify morphologically-distinct cell states associated with stemness, and determined the rates of transition to and from these states. These results demonstrate the utility of UPSIDE as a tool for unbiased exploration of cellular states and their dynamics from large, time-resolved imaging datasets.

## RESULTS

### Description of the UPSIDE platform

UPSIDE is designed to be a versatile machine-learning pipeline for unsupervised exploration cell morphological states in transmitted light images, and subsequent elucidation of their transition dynamics from movies (Figure 1A, see *Methods* section for detailed description of the pipeline). In this pipeline, cells are first segmented using a neural network that converts brightfield images of unlabeled cells into synthetic fluorescent images of cytoplasm for segmentation^21^. This neural network is trained using a set of images of cells stained for their cytoplasm (Supplementary Figure 1). This approach allows the network to autonomously tailor its parameters, and to accommodate a wide range of different cell types in order to optimize performance without human input. Dead cells and other debris are eliminated from identified cell sub-images through a convolutional classifier that is trained on manually annotated brightfield cell crops labeled as live or dead (Supplementary Figure 2).

**Figure 1.**
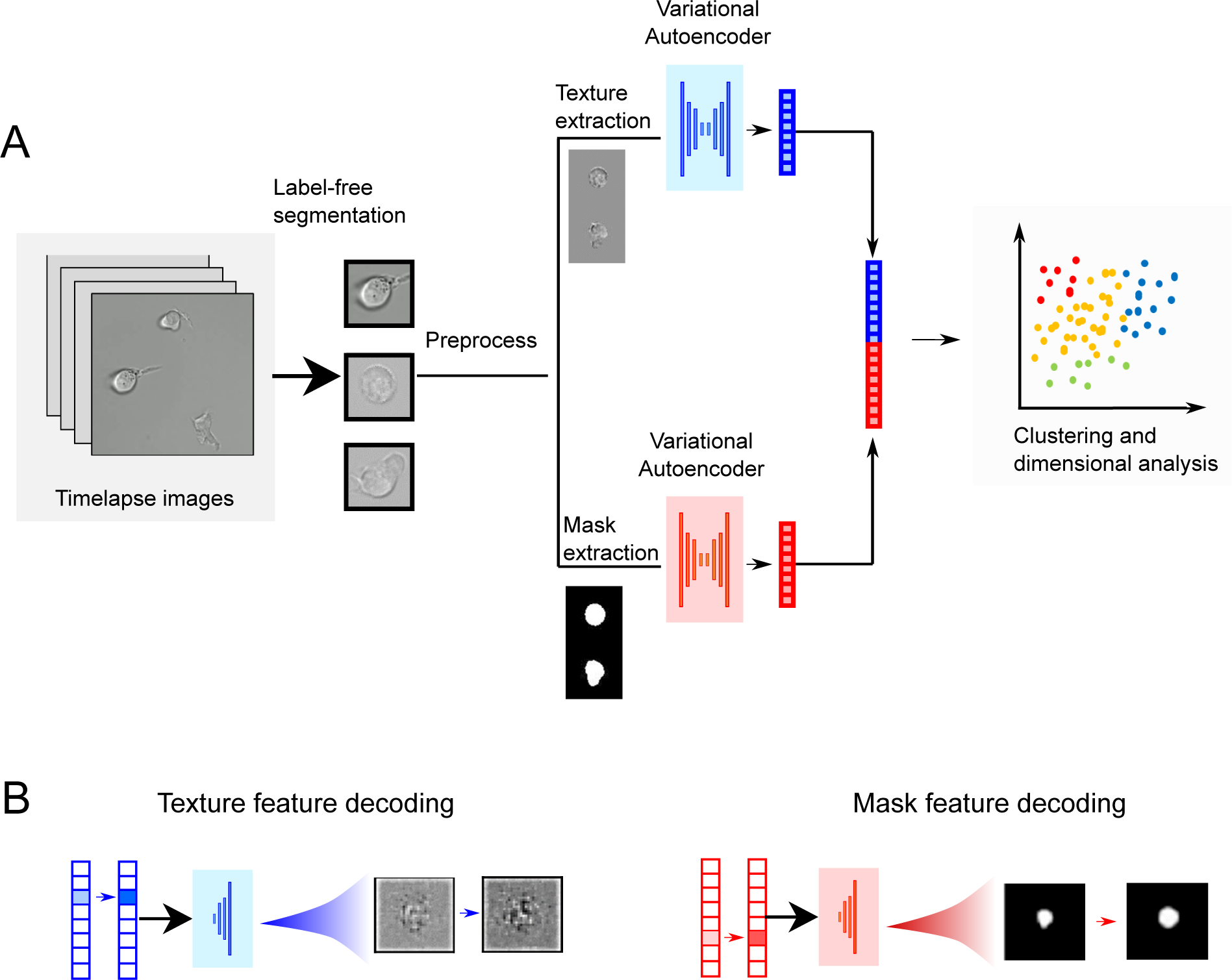
Description of the UPSIDE workflow. (A) Single cells are segmented directly from brightfield images and deep learning UNET architecture to predict synthetic fluorescent images^21^. Segmented cells are then pre-processed to generate separate mask and texture images, which are then used to concurrently train two variational autoencoders (VAEs). The shape and texture encodings learnt by these two VAEs are then concatenated and used for downstream data analysis. (B) Encoded latent vectors are then decoded into a shape and texture image to aid the interpretation of the encoded features.

UPSIDE utilizes a variational autoencoder (VAE) architecture to learn morphological features of segmented cells. Preprocessed masks of the cell and the cellular texture inside the mask boundary are then used to train two concurrent VAEs, one that encodes the cell shape, and another that encodes cell texture, through a binary mask. The mask VAE learns features related to overall cell shape and size, while the texture VAE learns pixel value variation within the mask itself while accounting for size and shape to some degree. Latent space encodings representing the learned mask and texture features are then multiplied with varying contributing weights, then concatenated for subsequent clustering and dimensionality reduction. Specifically, encodings are clustered into groups using the Louvain method^23^, represented on a 2D plane using the uniform manifold and projection algorithm (UMAP)^22^. Additionally, mask and texture vectors are subject to decoding, through variation of magnitudes of specific features or groups of features, followed by generation of synthetic images in observable image space (Figure 1B). This approach allows latent features to be visually displayed for human inspection and interpretation.

### UPSIDE distinguishes between different blood cell types in a heterogeneous population

To test UPSIDE’s ability to learn cell type-defining morphological features, we first determined whether this pipeline could distinguish among different cell types based on their morphologies in a mixed cell dataset. We focused first on using UPSIDE to discriminate among four blood cell types that, despite having distinct size, shape and textural features, were similar in their gross morphologies (Figure 2A, Supplementary Figure 3A): a mouse T cell leukemia line (Scid.ADH2), a mouse macrophage cell line (Raw246.7), a human acute myeloid leukemia cell line (Kasumi-1), and primary patient-derived human acute myeloid leukemia stem cells (CD34^+^CD38^-^ AML LSC). Brightfield images from each cell population were captured; cells were segmented using the neural network described above; and image crops of segmented cells were mixed together and encoded into the latent space using UPSIDE’s VAE (Supplementary Figure 4A). To quantify UPSIDE performance, we devised a cell type homogeneity score, which reflects how closely cells of the same type cluster together in their latent space (see *Methods* section). We ran the VAE for this dataset, optimizing combination weights between the learned mask and texture encodings to achieve the maximum mean homogeneity score across the four cell types (Supplementary Figure 5A-B and *Methods* section). To compare the performance of the VAE to other deep learning methods, we repeated this analysis with several alternative architectures such as a vanilla autoencoder (AE)^24^, the adversarial autoencoder with latent dimension encoding trained to fit a normal distribution or mixed gaussian distribution ^12^ (1xAAE and 4xAAE, respectively), and the ClusterGAN architecture ^25^ (ClusGAN) (see *Methods section*).

**Figure 2.**
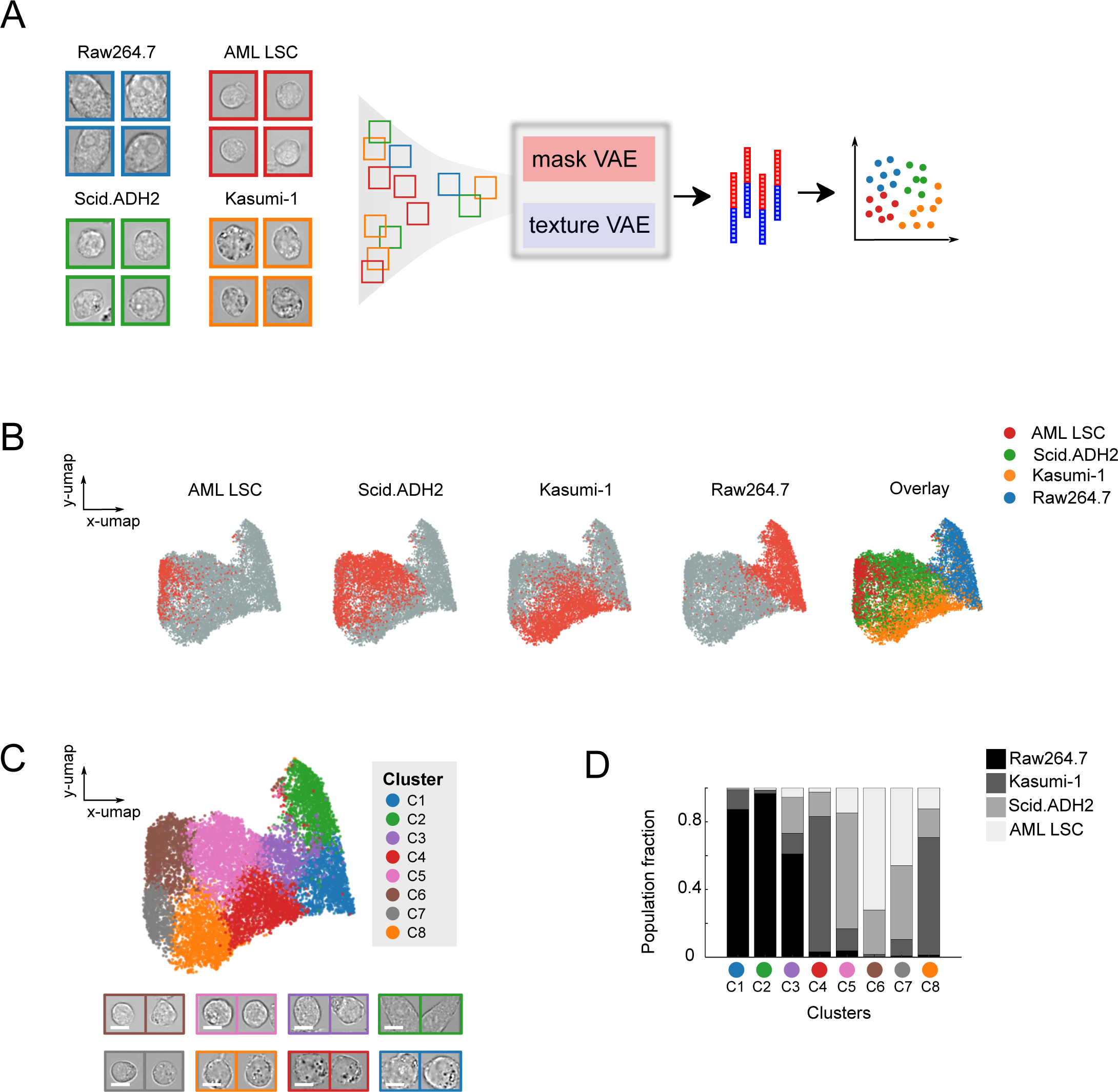
UPSIDE distinguishes morphologically-distinct blood cell types in a heterogeneous population. (A) Images of four different blood cell types were mixed together and passed through the UPSIDE workflow. Resultant shape and texture images were used to train concurrent VAEs. Output latent encodings were weighted relative to each other, concatenated, then projected onto a 2D plane using UMAP. (B) Dot plots show distribution of each cell type projected on 2D UMAP space made by UPSIDE. (C) 2D UMAP projection of the VAE-generated encodings that have been grouped into different morphological clusters using Louvain clustering algorithm. Representative brightfield cell crop images from the different clusters were listed. Scale bar represents 5 μm. (D) Cell type fractional composition within each cluster. A fixed number of cells from each cell type were sampled, and the cluster-wised cell type composition was calculated from this pooled population.

Our VAE outperformed these other approaches, generating approximately 6% higher homogeneity scores compared to the adversarial autoencoders, 9% higher compared to PCA, and 26% higher than the ClusterGAN architecture (Supplemental Figure 5C). Adversarial autoencoders performed better than the vanilla encoder, though worse than the VAE, possibly because it is difficult to train the discriminator to perfectly fit the latent encoding to the desired distribution. Surprisingly, ClusGAN architecture performed the worst, even though it generated quite realistic-looking generated cell texture and mask images (data not shown). This weaker performance may stem from an inability to consistently generate direct, regularized encoded representations. These comparisons suggest that the VAE architecture is particularly well suited for learning morphological features forcell type discrimination.

To further visualize and analyze the representation of cells in latent space, we projected the encodings from the VAE into two dimensions using the UMAP algorithm^22^ (Figure 2B). From the UMAP projection, we found that the cell types largely segregated into distinct regions in this two-dimensional space (Figure 2B). Raw264.7 macrophages occupied a region that was largely distinct from regions occupied by other three cell types, reflecting their markedly different cell size and shape distribution. The three other cell types occupied partially overlapping regions, reflecting greater similarities in morphology among these cells (Supplemental Figure 3A).

Interestingly, primary human AML stem cells (identified by their CD34^+^CD38^-^ surface marker phenotype) overlapped parts of the Scid.ADH2 region, suggesting some of Scid.ADH2 cells look quite similar to their AML counterparts. Despite these overlaps, there are substantial areas in the two-dimensional space occupied by these regions containing only one cell type, indicating the presence of morphological features that distinguish each of these three cell types from another and allow them to be identified in mixed populations.

To better understand the morphological features that drive cell type discrimination in this learned latent space, we clustered cell representations in the latent space using the Louvain method, then visualized cells and the morphological attributes that defined each cluster. Eight clusters were identified, with each enriched for different cell types (Figure 2C-D, Supplemental Figure 3B). Clusters C1-3 were highly enriched for Raw264.7 macrophages that are larger phagocytic cells than their progenitor cells. Clusters C4 and C8 were highly enriched for Kasumi-1 cells that contained circular profile cells with dark granules, a unique distinguishing observable feature of these cells. Cluster C5 was enriched for Scid.ADH2 cells, which were also circular, but lacked granules. Clusters C6 and C7 were enriched for both LSCs and Scid.ADH2 cells, both of which were small and lacked granules. Cells in Cluster C7 had darker interiors and less well-defined cell boundaries when compared with Cluster C6 cells, which indicate they are flatter and may be more substrate-adherent. The morphological differences within these clusters indicate the existence of distinct morphological sub-states within individual cell types.

To understand the morphological features that separate cells into distinct groups in latent space, we performed hierarchical clustering on the averaged latent space representation for cells from different clusters (Figure 3A). From this analysis, we found that each morphological cluster of cells was associated with a specific set of latent features, with magnitudes that are higher than population average. To decode these latent features, we transformed them back into synthetic images in visual space (Figure 3B-C, top). First, we generated a mean mask or texture vector by averaging over all cells in the dataset. From these mean vectors, we then selectively increased the magnitudes of the feature (or groups of features) of interest to generate a new vector. Using the VAE decoder module, we then transformed the feature-dominated vector and the mean vector into synthetic images for interpretation.

**Figure 3.**
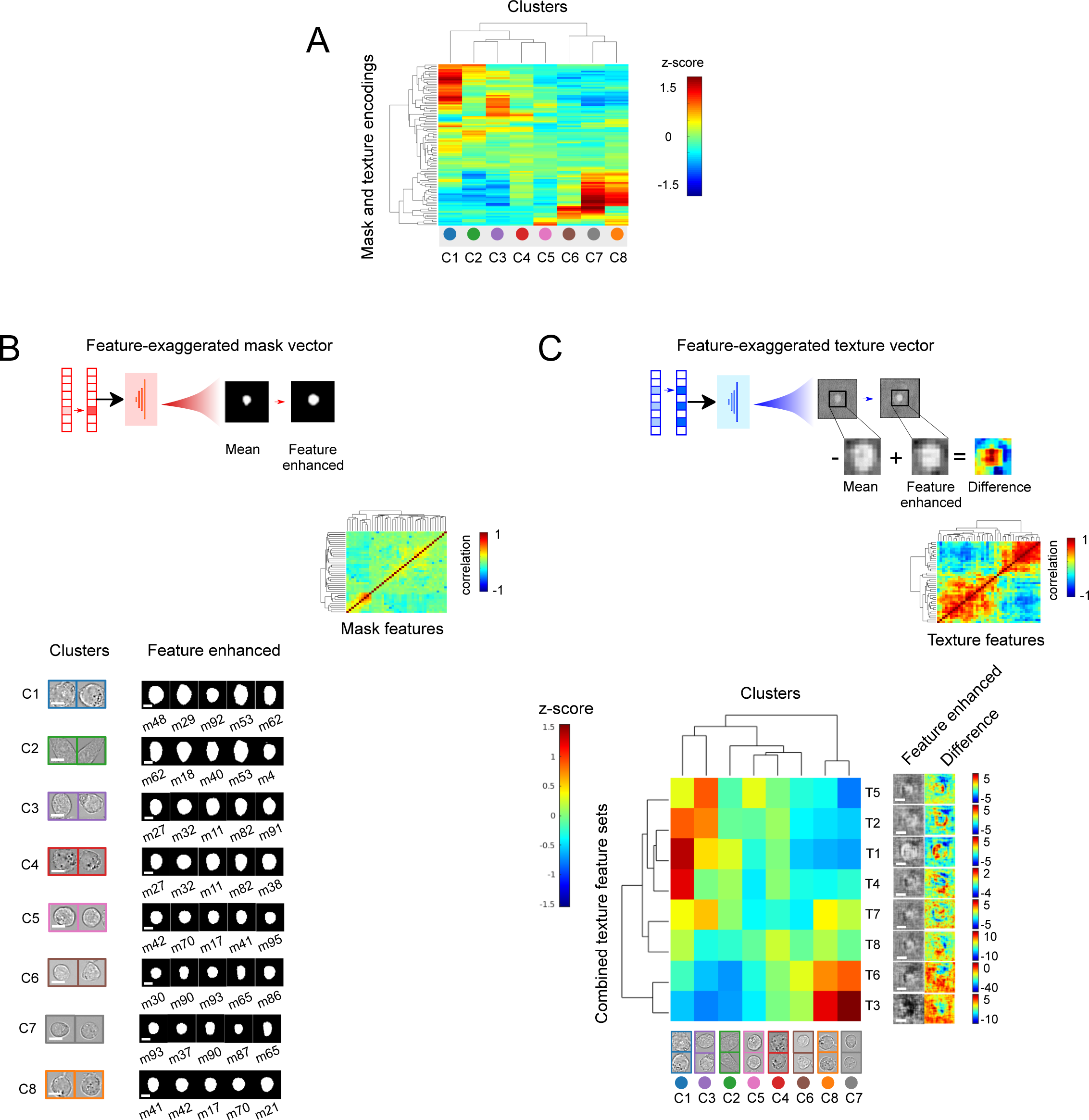
Cell-type specific morphological features can be interpreted by decoding the latent space cell representation. (A) Clustergram of average z-scores for latent shape and texture features for different cell clusters (see *Methods* section for how z-scores values were calculated). (B) Five mask features with highest z-scores for each morphological cluster are decoded and visualized. Inset: Clustergram shows matrix of correlation coefficients for forty mask features having the highest standard variation in the dataset. Scale bar represents 5 μm. (C) Individual texture features were clustered into eight groups (T1-T8) according to their correlation with each other (inset and clustergram). Each group was decoded into brightfield difference images for interpretation (see Methods). Scale bar represents 5 μm.

We first examined the synthetic decoded images from the five most enriched mask features for each morphology-defined cluster (Figure 3B, bottom). Clusters C1-4 contained large cells with large, round profiles. As expected, cluster C2 contained elongated cells with numerous elongated mask features. Clusters C5-8, in contrast, contained smaller cells enriched in features representing small, round profile cell shapes. These mask features are in general agreement with sizes and shapes for cells found within individual clusters (Figure 2C).

Unlike mask latent features, individual texture features in the latent space were not readily interpretable for this dataset (data not shown). However, because magnitudes of the projections along individual texture dimensions correlated strongly with each other in distinctive groups (Figure 3 C insets), in contrast to those for individual mask dimensions (Figure 3B, inset), observable texture features in the image space might be encoded not by individual latent features, but by groups of correlated latent features. Therefore, to visualize these observable features, we generated feature-dominated vectors by concurrently increasing groups of correlated latent features. We also calculated images representing the difference between feature-exaggerated decoded images compared to the mean texture decoded image for better visualization. From these synthetic difference images, we observed two overall texture pattern groups: one with darker cell interiors (T3, T6), indicative of a flatter morphology; and the other with lighter cell edges (T1, T2, T4, T5), indicative of a less flattened morphology (Figure 3C). The darker cell interior feature group is strongly enriched in Clusters C7 and C8, while the lighter cell edge group is significantly present in Clusters C1 and C3. Clusters C2, C4, C5, and C6 appear to have moderate enrichment in all these groups.

Taken together, these features reveal how UPSIDE separates cells into distinct morphological clusters by their size, shape and distinct textural features. This ability can be seen readily in Clusters C3 and C4, where cells of similar size and profile can be discriminated based on their cell edge texture features. Cells with similar textural features can also be discriminated using other features, e.g., Custers C7 and C8 are both enriched with dark cell interior textures, but differ in size with Cluster C7 cells larger on average than those in Cluster C8. These results demonstrate that UPSIDE can generate meaningful learned morphological features in an unsupervised manner, and these features can be effectively decoded into images to aid interpretability. This ability allows UPSIDE to extract valuable morphological properties by simply observing cells over time without prior manipulation or human annotations.

### UPSIDE uncovers morphologically distinct cell states in patient-derived leukemic cells

LSCs play critical roles in AML disease propagation and drug resistance^26, 27^. LSC and other AML cell subpopulations are typically identified and characterized by a combination of cell staining for granule content and cell surface markers as well as by their gene expression signatures^28, 29^. All of these classification approaches can be further extended by transmitted light imaging and analyses to provide complementary information about leukemic cell types and states that is not readily obtainable through more conventional classification approaches. In particular, live cell movies that resolve phenotypic states over time and in response to pharmacological treatment could provide unique insights into cellular heterogeneity and responses that could better inform therapeutic decision-making.

Towards this end, we employed UPSIDE to profile human LSCs cultured under cytokine conditions promoting expansion and differentiation, and filmed using brightfield imaging (Figure 4A, left). We directly isolated leukemic stem cells from an adult AML patient using as markers of stemness expression of the cell surface marker CD34 together with an absence of the differentiation marker CD38^26, 30, 31^ (i.e., the CD34^+^CD38^-^ cell fraction). To profile the self-renewal and differentiation dynamics of these sorted cells, we then cultured LSCs with either IL6 and thrombopoietin (TPO) to induce, to induce differentiation, or in the presence of Aryl hydrocarbon receptor inhibitors (AhRi) UM729 and SR1 to maintain stemness and suppress differentiation^32–34^. We then filmed these cells in the brightfield channel for ∼4 days at high temporal resolution (3 minute intervals, Figure 4A). To determine the association between observed cell morphological states, stemness and differentiation, we also added fluorescently-labeled anti-CD34 and anti-CD38 antibodies *in situ,* and took fluorescent images each hour to follow expression of these markers in imaged cells (Figure 4A, top right). Such *in situ* antibody labeling allows real-time visualization of cell surface marker protein expression with minimal effects on cell viability^35^. UPSIDE is well-suited to facilitate these types of time course analyses and image-based profiling: the use of brightfield imaging obviates the need for genetic engineering of fluorescent reporters to allow a wider range of analyses to be performed on primary patient-derived cell samples. Furthermore, a reliance on brightfield imaging minimizes cellular phototoxicity, thus enabling long-term cell observation at high temporal resolution.

**Figure 4.**
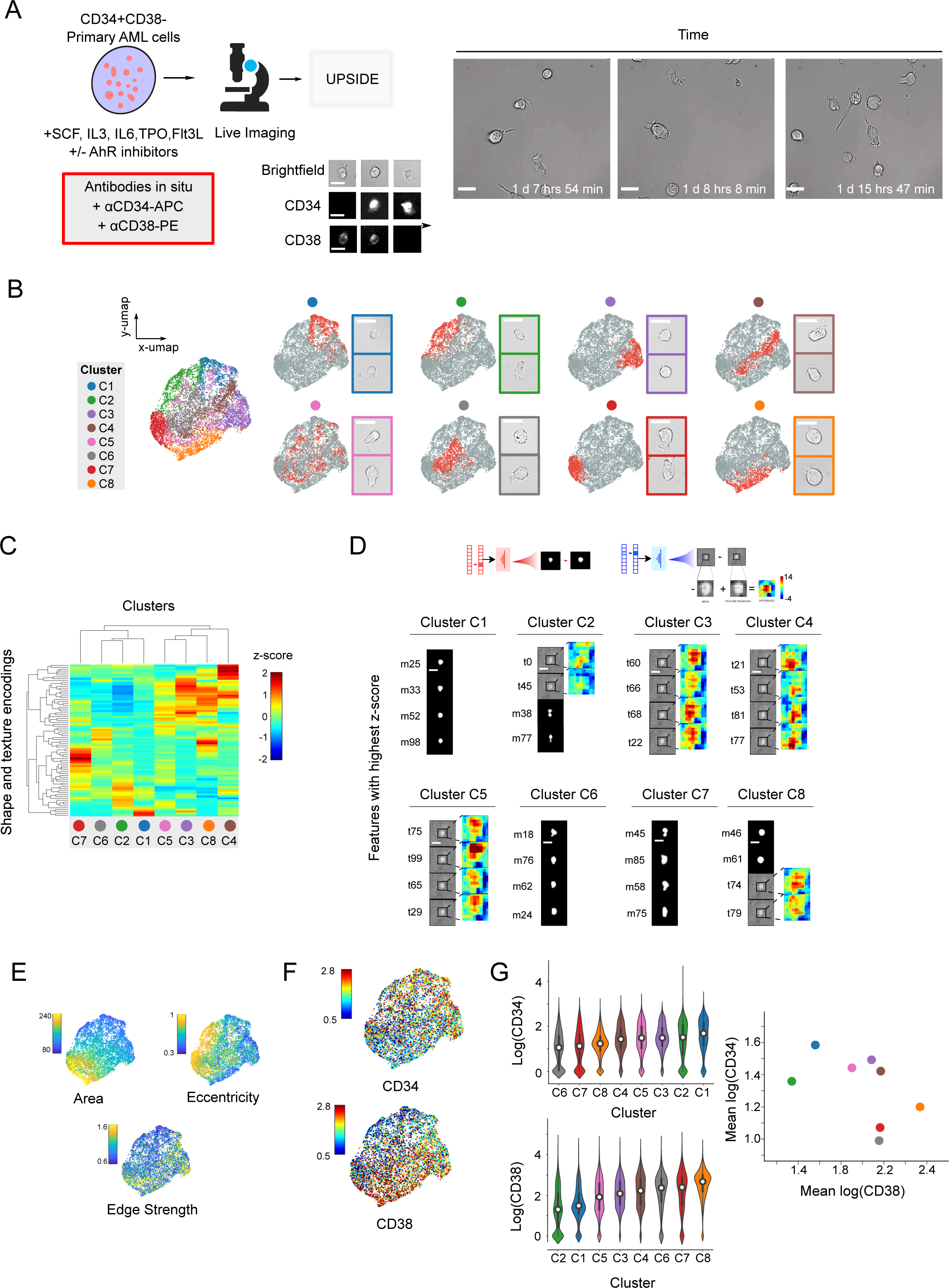
UPSIDE identifies stem cell-associated morphological states from patient-derived AML leukemic cells. (A) LSCs (CD34^+^CD38^-^) from an acute myeloid leukemia patient were cultured in cytokines with or without AhR inhibitors (UM729 and StemRegnin1) filmed for ∼5 days (left). Brightfield images were captured once every 3-5 minutes. αCD34-APC and αCD38-PE antibodies were added in situ, and fluorescent images were captured once every hour (top right). Still frames show representative time lapse images of AML cells (bottom right). Scale bar represents 10 μm. (B) UMAP 2D projection of the UPSIDE generated latent space cell representations. Individual morphological clusters were identified using the Louvain Clustering algorithm, then grouped manually based on their proximity to each other in the latent space (See Supplementary Figure 6B). Representative cell images from each cluster were also shown. Scale bar represents 10 μm. (C) Clustergram shows Z-scores of the latent mask and texture encodings for each morphological cluster. (D) Decoded images of the four most enriched features for each morphological state. Texture features were visualized using difference maps that were zoomed in around the decoded cells. (E) Area, eccentricity, and edge strength for each cell were calculated and mapped to the UMAP latent space representation. (F) CD34 and CD38 levels were mapped onto the UMAP. (G) Violin plots show distributions of CD34 and CD38 expression levels in different morphological clusters (left). Scatterplot shows mean CD34 levels against CD38 levels for each morphological cluster (right).

From the resultant timelapse images, we observed considerable heterogeneity in the morphologies of observed cells, with these cells differing in their sizes, shapes and textures, as observed by their observed contrast from transmitted light images (Figure 4A, right, Supplementary Movie S1). To better understand the morphological states of these cells, we fed these images into the UPSIDE pipeline. Cells across all time points were segmented, then encoded using separate shape and texture VAEs (Supplementary Figure 4B). We then organized representations of cells in latent space into morphological groups using the Louvain clustering method (Supplementary Figure 6). Based on proximity in the latent space, we further combined some of these groups into larger clusters. To visualize obtained cell clusters in two-dimensional space, we then projected these cells’ latent representations onto a two-dimensional plane using the uniform manifold approximation and projection algorithm (UMAP). This projection revealed the locations of the discrete clusters, along with their overlap regions. In this two-dimensional visualization, some cell clusters showed considerable boundary overlap with others, likely reflecting the continuous nature of the latent features encoded by the VAE.

In order to gain insight into the features that drive the separation of the cell encodings into distinct clusters, we performed hierarchical clustering on averaged cell encodings from each group (Figure 4C), then decoded the specific mask or textural features with the highest z-scores in each group to generate feature-dominated synthetic images as described above (Figure 3B-C).

These synthetic images highlight significant morphological features that display coherence across cells within a cluster, but differ between cells in different clusters (Figure 4D, Supplementary Figures 7-8). Examples of coherent morphological features include size, with some having smaller cells (Clusters C1,C2) and others having larger cells (Clusters C6,C7,C8); cellular elongation or eccentricity, with some clusters displaying rounder cell profiles (Clusters C1,C8) and others have more elongated cells (Clusters C2 and C7); and the presence of finer morphological features, such as cytoplasmic protrusions from the cell body (Clusters C6 and C7). Another important coherent morphologic feature was cell edge and boundary morphology. Cluster C8 cells had pronounced cell edges all around the boundary, while other cells had dimmer, asymmetric cell edges (Clusters C4,C5), as visualized with synthetic difference images of cell texture. These features indicate cell flattening or protrusion out from the imaging plane in the Z-direction, and may reflect differences in cell adhesion to substrate (Supplementary Figure 7). These decoding results indicate that clusters identified by UPSIDE contain leukemic cells with distinct morphological states.

To verify that these differences in decoded features indeed reflect systematic morphological differences between cells in different clusters, we calculated cell area, eccentricity, and edge strength – defined by the maximum value of the cell’s gradient image – and then plotted these quantities onto the 2D projection of the latent space (Figure 4E). Indeed, regions occupied by the different cell clusters had area, eccentricity, and edge strength values consistent with what was generally observed in the decoded cell images: Clusters C1-3 resided in the region with small cell areas, whereas Clusters C6, C7 and C8 resided in the region with larger cell areas. Elongated cells in Clusters C2 and C7 resided in regions with high eccentricity, whereas cells with darker cell edges in Clusters C3 and C4 resided in regions with high edge strength. Together, this analysis shows that UPSIDE can elucidate defining characteristic shape and textural features of cells in different morphological states.

### Distinct morphological states are associated with different degrees of stemness

Cells in the different morphological states identified above may exhibit different degrees of AML cell stemness or differentiation. To test this idea, we quantified CD34 and CD38 expression levels for each cell, and mapped them onto the 2D projection of AML cell’s learned latent dimensions (Figure 4F-G). We then calculated average CD34 and CD38 levels for all cells in a cluster, and plotted these values against each other. These plots show that the different morphological cell clusters differ in their average CD34 and CD38 levels, indicating different degrees of cellular maturation. Cells in clusters C1 and C2 had high CD34 but low CD38 expression. Cells in these clusters differed in their degree of elongation, but were mostly smaller and flat, indicating that the most stem-like cells that are characterized by their small size and their flatness, possibly because of strong substrate-attachment through the expression of cell adhesion proteins^36^. Cells in Clusters C3, C4 and C5, which were larger and appear less flattened, had high expression of both CD34 and CD38, indicating that they may reside in an intermediate maturation state. Finally, cells in Clusters C6, C7 and C8, which were uniformly larger but had varying degrees of flatness, had low CD34 but high CD38 expression. Thus more mature cells may be generally larger, despite continuing to display variable shapes and degrees of substrate adherence. These results highlight the ability of UPSIDE to discover subtle yet biologically relevant cell morphological states in heterogeneous populations.

### Population dynamics of cell morphological states

To gain insight into the population dynamics of cells in different morphological states, we examined how the numbers of cells in different clusters evolved over time, both with and without suppressing differentiation with AhR inhibitors (AhRi; Figure 5A, Supplementary Figure 9A) .

**Figure 5.**
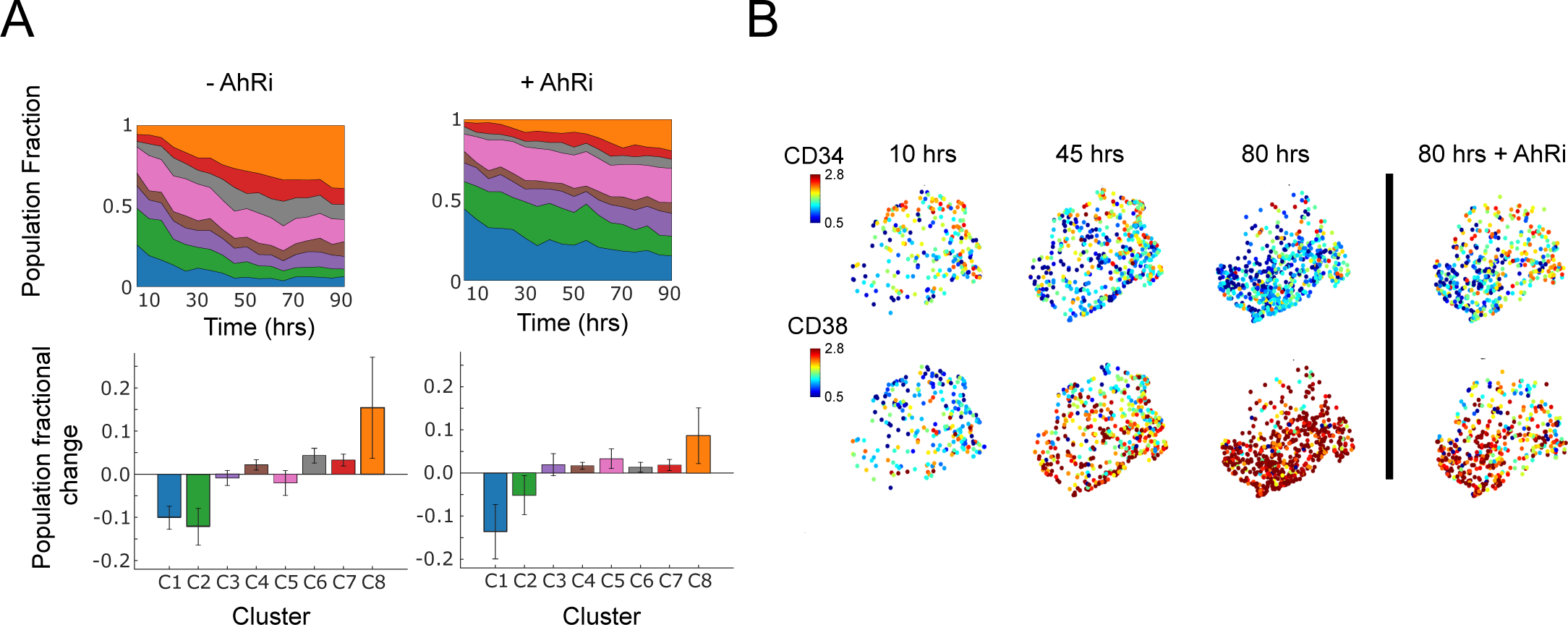
Population dynamics of identified morphological states. (A) Population fraction dynamics over time for each morphological cluster with (right) or without (left) AhR inhibitors (top). Population fraction difference for each cluster between the first 20 hrs and last 10-20 hrs of culture (bottom). (B) UMAP showing CD34 and CD38 expression levels at different time points, in the presence or absence of AhRi.

In the absence of AhRi, cells with stem-like morphological states (Clusters C1 and C2) were progressively depleted, while those with differentiation-associated morphological states (Cluster C8) expanded, consistent with the maturation of LSCs into more differentiated cells over time. In the presence of AhRi, there was a less pronounced increase in differentiated morphological states with a concomitant decrease in stem-like states. This reflects the known effects of AhRi in maintaining stem cell self-renewal. Interestingly, the fractions of cells in intermediate morphological states (Clusters C3-7) remained largely unchanged, regardless of the presence of AhRi, suggesting that AhR inhibition may affect intermediate state transitions without driving specific outcomes. At the same time, CD34 levels decreased whereas CD38 levels increased over time, with both these changes becoming less pronounced with the addition of AhRi (Figure 5B, Supplementary Figure 9B). Together, these results provide insights into the kinetics of LSC self-renewal and differentiation, and how these kinetics are affected by pharmacological compounds that modulate self-renewal.

Parallel experiments were used to further explore morphological changes that reflect LSCs maturation. We cultured LSCs (CD34^+^CD38^-^) in parallel with live imaging experiments, and analyzed them after three days for expression of CD34, CD38, and CD123, another common LSC marker^37^ (Supplementary Figure 9 C). Compared to untreated samples, cells treated with AhRi show higher expression of CD34 and CD123. On the other hand, CD38 expression magnitude was higher in the untreated sample, indicating greater differentiation in this population. Of note, a population of cells expressed both CD38 and CD34; this result indicates that expression of these markers is not mutually exclusive. Parallel live imaging experiments of AhRi-treated cells showed slower expansion of large round cell morphology clusters compared to the untreated counterpart (Figure 5A, Supplementary Figure 9A). Together, these results demonstrate that distinct cell morphological states identified using UPSIDE indeed reflect leukemic cells in different states of maturation.

### Inference of morphological state transition rates by cell linkage analysis

The high temporal resolution of the brightfield movie analyzed above enables tracking of individual cells from frame-to-frame and, in conjunction with UPSIDE, inference of the rates at which leukemic cells transition between different morphological states. Here, we develop an analysis routine to automatically infer transition rates from brightfield movies. In particular, this UPSIDE-enabled analysis obviates the need for the generating individual cell tracks, which are typically error prone and requires considerable manual intervention. We paired cells from adjacent frames together, based on their close proximity. We then identified the morphological states of linked cell pairs using the VAE above (see *Methods* section), and calculated state transition probabilities based on the frequencies of linked cell pairs with specific initial and final morphological states (Figure 6A). By repeating this calculation over all possible pairs of morphological states, we obtained a matrix, describing the transition probabilities between different morphological states (Figure 6B, left).

**Figure 6.**
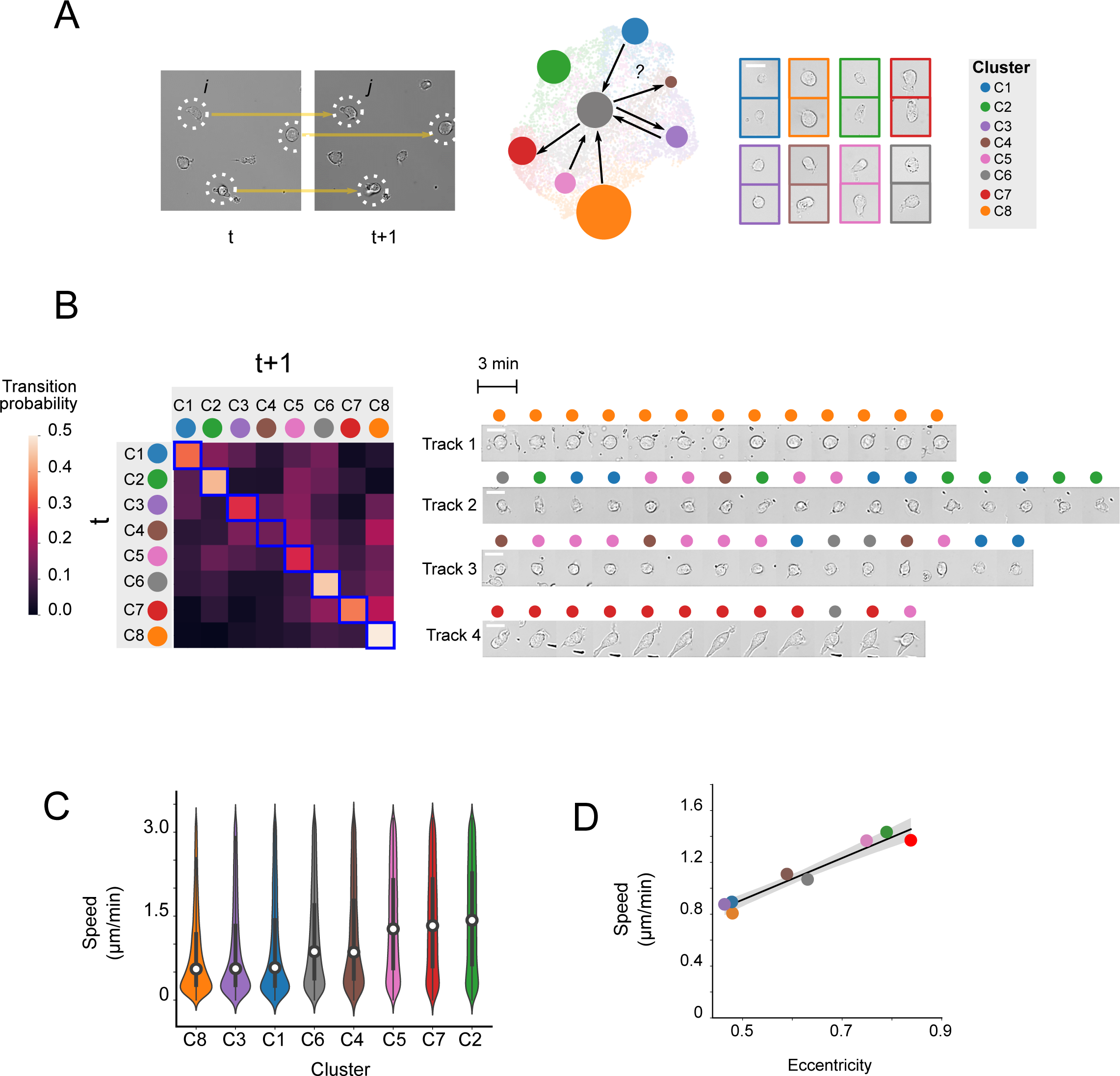
Calculation of morphological state transition probabilities by cell linkage analysis. (A) Cell pairs found in proximity across on successive time points were linked (left). Cell linkages, along with assigned morphological states of linked cells, were used to calculate transition probabilities between all states. (B) Heatmap shows transition probability matrix between all morphological clusters (left); image montage shows representative cell tracks identified from the culture (right). Scale bar represents 10 μm. (C) Distribution of cellular velocity for linked cells for each morphological cluster. (D) Plot shows mean cell velocity against mean cell eccentricity for each morphological cluster.

This analysis showed that different morphological states showed considerably different stability and transition preferences (Figure 6B, left). Some states represented by Clusters C2, C6, C7 and C8 had high stability, as indicated by a high probability of remaining in the same state from one frame to the next. These represent a variety of morphologies, from small, elongated cells to larger, rounder cells. Other states – such as clusters C3, C4 and C5 – showed lower probabilities of remaining in the same state, and higher probability of transitioning into a different state by the next frame. To verify these inferred transition rates, we visually generated and inspected a number of different single tracks for cells in different morphological states (Figure 6B, right).

Indeed, cells starting in states C7 and C8 (tracks 1 and 4) tended to remain in the same state, whereas cells starting in states C1, C3 and C5 were highly dynamic, switching from one state to another rapidly in successive frames (tracks 2 and 3). Notably, transitions occurred preferentially between particular groups of states. For instance, track 2 showed frequent transitions between states C1 and C2, whereas track 3 showed frequent transitions between states C4, C5 and C6, consistent with the elevated transition probabilities between these states as observed in the transition matrix (Figure 6B, left).

The morphological state transitions observed above occurred over tens of minutes (Figure 4B), a timescale much shorter than that for cell differentiation, which occurs progressively over the course of movie observation (Figure 5A). As such, these transitions are unlikely to report on an entire cell differentiation event, but rather snapshots of such a process. They could also reflect more rapid cell phenotypic state changes. Cells tend to polarize as they crawl on a substrate; thus, it is possible that some of these observed transitions could reflect transitions from a stationary to a motile state. To test this hypothesis, we derived the instantaneous velocities of cells in different morphological states, by calculating the displacement between successive frames for each state (Figure 6C). From this analysis, we found that cells with elongated morphologies, such as those in states C2, C4, C5, and C7, showed a higher movement velocity compared to other states. Consistently, there was a strong correlation between instantaneous velocity and cell eccentricity, averaged over all cells in individual clusters (Figure 6D). Thus, it is likely the case that the observed morphological transitions partially involve the rapid switching between stationary and mobile states (Figure 6B, left, tracks 2-3). Interestingly, both stem-like (CD34-high) and more differentiated (CD34-low) cells contained both stationary and moving cells populations. This indicates that cell mobility, or lack thereof, is not necessarily strongly tied to the differentiation process.

Cells in different morphological states differ not only in their degrees of elongation and, by extension, motility, but also in other morphological characteristics that are associated with stemness; thus, cell state transition probabilities would also likely depend on rates of differentiation. To test whether this was the case, we repeated this state transition analysis for cells treated with AhR differentiation inhibitors (Supplementary Figure 10A). In the presence of AhRi, there was a decrease in the transition probabilities into the more differentiated states (C6, C7 and C8) Together with an increase in the transition probabilities into and amongst intermediate cell states (C3, C4 and C5). This suggests that AhRi blocks the conversion from a stem-like morphological state to a differentiated state by stalling cells in an intermediate state. Single cell track analyses showed that cells often transitioned rapidly from an intermediate state (C5) to more mature state (C8) without AhRi, but stalled at an intermediate state (C5) when AhRi is present (Supplementary Figure 10B). Taken together, these observations show that changes in cell differentiation dynamics can influence the probabilities at which cells transition between different morphological states.

## Discussion

Here, we presented UPSIDE, a deep learning platform for unsupervised exploration of dynamic cell morphological states in transmitted light movies. Using UPSIDE, we identified cell morphological features and states in a heterogeneous collection of blood cell types. We found that the UPSIDE VAE learning architecture outperforms other comparable methods in recognizing unique morphological features within each cell type. We further demonstrated the utility of our method by uncovering morphological states in primary human AML patient-derived leukemic cells displaying different degrees of stemness, differentiation, and cellular mobility. Finally, UPSIDE addresses the issue of latent feature interpretability, one of the most challenging aspects of analyzing deep convolutional networks, to allow more intuitive insight into learned latent morphological features.

UPSIDE provides a versatile new platform to enable the use of timelapse live imaging to discover phenotypic states and analyze their dynamics in heterogeneous cell populations. This method does not require genetic manipulation for labeling, identifying or tracking cells, and thus enables observation and analysis of primary cells such as patient samples. The lack of reliance on fluorescence imaging reduces phototoxicity burden, which may have the added advantage of allowing imaging at higher time resolution over long times, thus allowing dynamics analyses over a wide range of timescales. Finally, UPSIDE is an unsupervised method which tailors analyses and outputs to differences within each individual dataset. Thus no initial knowledge of the system, feature selection or assumptions are necessary, as the VAE architecture is capable of self-learning the distinguishing features.

The ability of UPSIDE to resolve the dynamics of state transitions opens doors for systematic analysis of cell response dynamics in primary cells and other genetically intractable cell types. In particular, one attractive application for this pipeline is for automated, high throughput analysis of leukemic cell responses to chemotherapy. AML drug resistance poses a significant clinical challenge as the majority of patients eventually develop relapse disease. Growing evidence suggests leukemic stem cell (LSC) populations in AML harbor drug-tolerant sub-populations that survive drug treatment and eventually lead to relapse disease^30, 38^. To this end, a large scale screen of hundreds of drug treatment or regimen time courses can be captured via brightfield time lapse imaging^39, 40^. UPSIDE can then be employed as an unbiased method to survey and specify important morphological features associating with therapeutic response, persistence or resistance as a function of cell types, cell states and treatment. We anticipate UPSIDE will provide a valuable tool to enable the broader application of powerful deep learning to a wide range of biological questions.

## METHODS

### Experimental Techniques

#### Cell Lines

Kasumi-1, Scid.ADH2, and RAW246.7 cell lines were cultured Eagle’s minimal essential medium (DMEM), phenol-red free containing 10% Fetal Bovine Serum (FBS), Penicillin-Streptomycin-Glutamine (Gibco 10378016) at 37°C and 5% CO_2_ (ThermoFisher) for 2 days before imaging. For 5 cell type imaging experiments, each cell line was imaged in a separate individual well in the same 96-well glass-bottomed plate. AML211 CD34+CD38- subpopulation was cultured in ‘Differentiation Media Condition’ for 2 days before imaging at the same time with the cell lines.

#### Culture of patient-derived leukemic cells

Primary acute myeloid leukemia samples (AML211) were provided by the Pamela Becker lab. The study was conducted with approval of the Institutional Review Board, Fred Hutchinson Cancer Research Center. The samples were obtained from AML patients with informed consent.

Cryo-preserved AML cells were thawed in ‘Long Term Bone Marrow (LTBM) Media’ [Iscove’s Modified Dulbecco’s Medium (IMDM) with glutamine and HEPES (Mediatech. Inc, Manassas, VA), 15% Fetal Bovine Serum (HyClone, Logan, UT), 15% Horse Serum (VWR), 50 μM beta-mercaptoethanol (Sigma), 0.043% Monothiolglycerol (Sigma)] and washed twice to remove DMSO, then cultured in LTBM with 50U/ml DNase to break up and free live cells if chunks are present at 37°C and 5% CO_2_ for 1 hour. Cells were then cultured in LTBM with 10 ng/ml recombinant human Stem Cell factor (SCF) at 37°C for 2 days. For cell sorting and flow cytometry analysis, cells underwent Lymphocyte Separation Medium (Mediatech. Inc, Manassas, VA) to remove dead cells and were stained with CD34 (ThermoFisher 17-0349-42), CD38 (ThermoFisher 12-0388-41), and CD45 (VWR 10758-692) for flow cytometry analysis and sorting by FACS Aria (BD biosciences, San Jose, CA).

Sorted CD45+CD34+CD38- subpopulation from AML211 samples were cultured in ‘Differentiation Media Condition’ (adapted from (Klco et al., 2013)) which comprises of Eagle’s minimal essential medium (DMEM), phenol-red free containing 10% FBS, Penicillin-Streptomycin-Glutamine (Gibco 10378016), 100 ng/ml Recombinant Murine SCF (Prepotech 250-03), 50 μM beta-mercaptoethanol (Sigma M6250), 10 ng/ml Recombinant Human IL-3 (Prepotech 200-03), 20 ng/ml Recombinant Human IL-6 (Prepotech 200-06), 10 ng/ml Recombinant Human TPO (Prepotech 300-18), 10 ng/ml Recombinant Human Flt3-Ligand (Prepotech 300-19), or ‘Maintenance Media Condition’ (adapted from (Pabst et al., 2014)) which comprises of minimal essential medium (DMEM), phenol-red free containing 10% FBS, Penicillin-Streptomycin-Glutamine (Gibco 10378016), 100 ng/ml Recombinant Murine SCF (Prepotech 250-03), 50 μM beta-mercaptoethanol (Sigma M6250), 20 ng/ml Recombinant Human IL-3 (Prepotech 200-03),), 50 ng/ml Recombinant Human Flt3-Ligand (Prepotech 300-19), 1 μM UM729 (STEMCELL Technologies 72332), and 500 nM StemRegenin-1 (STEMCELL Technologies 72342). Cells were cultured on treated polystyrene (Corning) of glass-bottomed (Mattek) 96-well culture plate coated overnight with 33.33 μg/ml Retronectin (Takara T202).

For imaging differentiation assay, CD34 Human Monoclonal Antibody (4H11), APC (eBioscience 17-0349-42) and CD38 Human Monoclonal Antibody (HB7), PE (eBioscience 12-0388-41) was spiked into the culture media. Cells were imaged every 3-5 min with brightfield and 60 min with fluorescent light for 4 days.

#### Image Acquisition

Timelapse imaging was performed on Inverted Microscope Platform Leica DMi8 (Leica Microsystem). All image acquisitions were performed using 40X air objective.

Fluorescent images were captured using Laser Diode Illuminator, LDI (89 North).

### Image Analysis

UPSIDE computational pipeline is designed to analyze the morphological diversity of cells from timelapse brightfield images. The method consists of four main modules: 1) Label-free prediction, 2) Image segmentation, 3) Live cell classification, and 4) Unsupervised feature learning. The following section describes each of the modules in further details.

### Label-free Imaging and Image Segmentation

UPSIDE utilizes a label-free imaging method to identify cells from brightfield (BF) images. Here we adapted a U-net-based deep learning technique described by ^21^ to predict fluorescent pictures of the cells’ cytoplasm and nuclei from the captured BF images. To complete this task, a small sample of cells to be analyzed was stained CellTrace Violet Cell Proliferation dye (ThermoFisher C34557) to label the cytoplasm.

Training data was obtained by capturing approximately 300 – 400 BF images and their corresponding nuclei or cytoplasm fluorescent images. This data was then used to train a model that predicts nuclei and another that predicts cellular cytoplasm. These two models were subsequently used to predict fluorescent images for the main timelapse brightfield image stacks. Object segmentation was performed on predicted nuclei images using ictrack software described in^19^ .

### Live Cell Classification

Identified cell crops were then fed through a classifier that separated a living cell for further analysis from dead cells, and other debris that were picked up from the segmentation. We built a convolutional neural network for this classification task:

To obtain training data for this network, ∼10000 brightfield cell crops were manually annotated as ‘Live’ or ‘Dead.’ The network were trained for approximately 10,000 steps, and cross entropy loss were calculated a Adam optimizer^41^ were used for weights and biases learning:

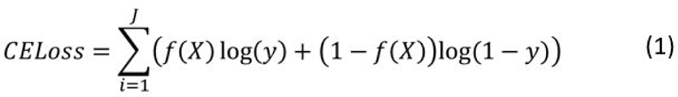

Where *f* (*X*) is the predicted class of a given cell crop *X* and *y* is its correct label. The remainder of the identified cell crops were then fed to the trained classification model. Crops classified as ‘Dead’ were discarded, and ‘Live’ crops were used for further analysis.

### Unsupervised Feature Learning

Morphological feature learning in UPSIDE relies on the variational autoencoder architecture (VAE) ^20^ to perform feature extraction. Two information pieces were used to train the VAE: 1) The overall shape of the cell and 2) The cellular texture inside the boundary mask. Predicted CellTrace violet signal of the cell was used to generate the cell shape crop. The following image preprocessing steps were performed to minimize trivial variations between cell crops:

- Object re-centering
- Object rotational orientation to 90°. All cell crop images are then rescaled accordingly to eliminate image’s dimension inflation due to rotation.
- Object’s vertical and horizontal pixel density reorientation to top and right, respectively

To obtain texture representation, brightfield pixel values distribution inside the cell’s mask was scale adjusted to zero mean and unit variation. They are then scaled linearly to be between 0 and 1 to facilitate learning with VAE. All pixel values outside the boundary were set to 0.5.

Preprocessed image crops for shape and texture were used to train two separate VAEs. The overall architecture is as described below:

The loss function for the VAE is a weighted combination between reconstruction loss and Kullback-Leibler Divergence loss:

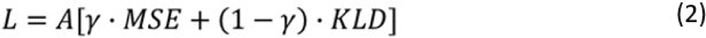

Where A is a constant, and γ is between 0 and 1. Additionally,

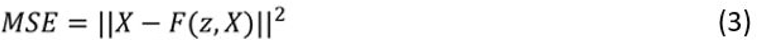

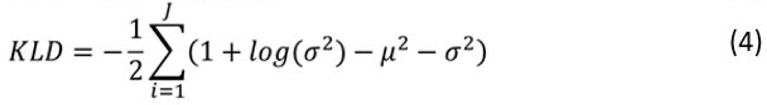

VAE for cell shape feature extraction was first trained for ∼100000 steps while VAE for texture feature extraction was first trained for ∼200000 steps. Trained weights and biases for the cell shape extraction were then used to encode all cell crops obtained from the movie into 100-element vectors. These vectors were projected onto a 2D plane using UMAP ^22^. Cell crops with defective shapes are gated out using the cytometry2 function in ictrack. The remaining crops were then used to train VAE for cell shapes and texture separately. Afterward, cell crops were encoded into 100-element shape vectors and 100-element texture vectors. Each cell crop’s latent vector is represented by a weighted concatenation between the shape and the texture contributions:

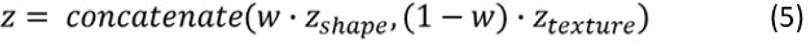

Encoded latent dimension of cell crops are then clustered using Louvain clustering algorithm. To generate synthetic images, encoded cell barcodes and arithmetic variations were treated as z and fed directly into the decoder.

### Comparable Deep Learning Architectures

In addition to utilizing the Variational Auto Encoder architecture to learn the latent dimensions in our imaging datasets, we tested a few other deep learning architectures to compare their performances with our current approach:

#### Vanilla Auto Encoder (AE)^24^

In this architecture, each processed shape or texture is fed through a series of convolutional layers and fully connected neural network layers to generate a latent vector with a dimension of 100. The organization of the neural network layers are as follows:

The loss function for the AE is:

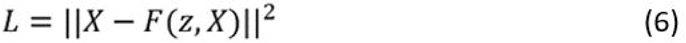

AE for cell shape feature extraction was first trained for ∼100000 steps while AE for texture feature extraction was first trained for ∼200000 steps. Each cell crop’s latent vector is represented by a weighted concatenation between the shape and the texture contributions:

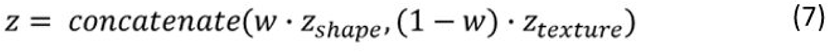

#### Adversarial Auto Encoder (AAE) ^12^

In this architecture, each processed shape or texture is fed through a series of convolutional layers and fully connected neural network layers to generate a latent vector with a dimension of 100. The latent dimension was then regularized using a discriminator that forces the dimension space into a unit gaussian distribution (1x AAE) or four mixed gaussian distributions (4x AAE). The organization of the neural network layers are as follows:

The loss functions for the VAE are:

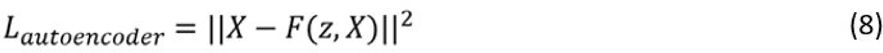

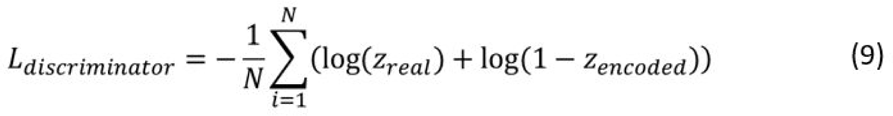

Where *z _real_* is a 100 element vector sampled from a normal gaussian distribution (1X AAE) or a mixed 4-gaussian distribution with each gaussian’s mean to be -1, -0.5, 0.5, and 0.5 and standard deviation to be 1 (4X AAE).

AAE for cell shape feature extraction was first trained for ∼100000 steps while AAE for texture feature extraction was first trained for ∼200000 steps. Each cell crop’s latent vector is represented by a weighted concatenation between the shape and the texture contributions:

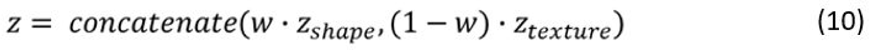

#### *ClusterGAN*^25^

This architecture carries an encoder that converts a generated image into a latent dimension which is then forced to match the same starting latent code that was originally used to make the image. This is a semi-supervised architecture where a specific number of classes needs to be predetermined beforehand. To convert this into an unsupervised method, we removed the class module, enabling the GAN to draw data from a normal distribution, without the one-hot class vector input. The neural network organizations for the generator, encoder, and the discriminator are follows:

Loss functions used for training were described previously ^25^. We input the cell crops into the encoder module of ClusterGAN to generate the latent dimensions for the comparative analysis with other architectures.

Cell shape feature extraction was first trained for ∼100000 steps while the texture feature extraction was first trained for ∼200000 steps. Each cell crop’s latent vector is represented by a weighted concatenation between the shape and the texture contributions:

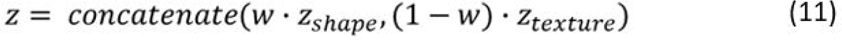

### Algorithms and quantitative analysis

#### Neighbor similarity scoring

The metric is formulated to estimate the degree of homogeneity of the grouping of each cell type in encoding space of the four cell types data. Specifically, the neighbor similarity score *H_C_* for a given cell type *C* is define as followed:

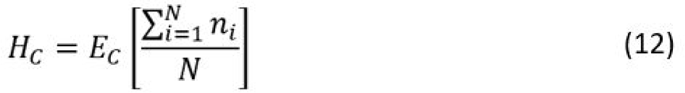

Where *E*(·) represents the expectation value, or the mean over all cells within a cell type *C*, and *N* specifies a predetermined number of nearest neighbor cells to a given cell. Furthermore, for each neighboring cell *i*, *n_i_* = 1 if the identity of *i* is *C*, and *n_i_* = 0 otherwise.

#### Latent dimension z-score calculation

The z-score *Z_f_* _,*c*_ of a particular feature *f* of cluster *C* is defined as the fold difference in standard deviation between the mean of the value of that feature in cluster *C* compared to that of the complete dataset:

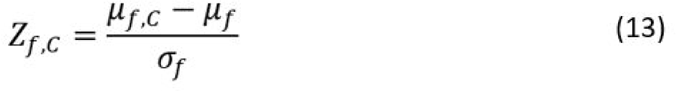

Here, μ*_f_* _,*C*_ is the mean value of feature f over all cells in cluster *C*, and μ*_f_*, σ*_f_* are the mean and standard deviation of feature *f* over the dataset.

#### Pairwise cell tracking algorithm

The pairwise cell tracking algorithm was built to ensure the validity of a given paired cell linkage from one frame to another. To achieve this goal, we established stringent requirements for a given cell pair to be considered ‘valid’. Specifically, the linking algorithm concerning all cells in frame t is as followed:

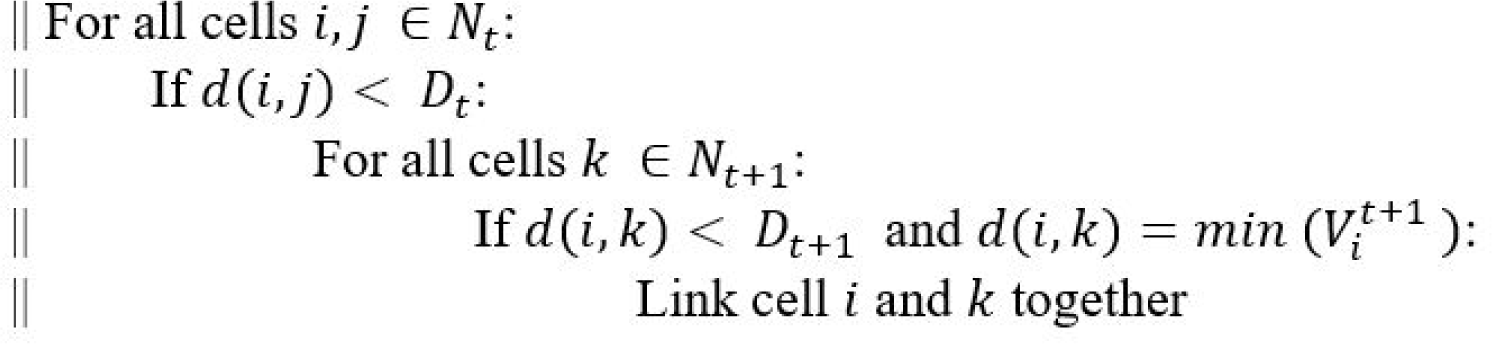

Here, *N_t_* represents a set of all detected cells in frame *t* ; *d*(*a*, *b*) denotes the Euclidean distance between cells *a* and *b*, and *V ^t^* represents a set of Euclidean distances between cell *a* in frame *t* − 1 to all cells in frame *t* .

#### Transition probability between cell clusters

In order to estimate the transitional dynamics between identified morphological clusters through time, we determine the transition probability between a given cell *X* at time *t* to belong to cluster *i* to another cluster *j* in a set of clusters *C* at time *t* + 1 as followed:

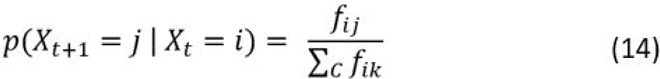

Where *f _ij_* is the number of transitions from *i* to *j* .

## Data Availability

Imaging data for the UPSIDE platform will be made available on the Image Data Resource (https://idr.openmicroscopy.org/about/). Code for the software is available on Github (https://github.com/KuehLabUW/UPSIDE).

## Acknowledgements

We thank Chek Ounkomol, Gregory Johnson, and Molly Maleckar for advice on label-free image segmentation and deep learning architecture design. We also thank members of the Kueh lab for discussions and feedback on the manuscript. This research was funded in part by an NIH Pathway to Independence Award 5R00HL119638 (to H.Y.K.), NIH/NHLBI grant R01HL146478 (to H.Y.K), a Tietze Foundation Stem Cell Scientist Award (to H.Y.K.), the NIH/NCI Cancer Center Support Grant P30CA015704 (to H.Y.K. and P.S.B.); and NIH/NCI Award P01CA077852 (to R.M.). The content is solely the responsibility of the authors, and does not necessarily represent the official views of the National Institutes of Health.

---

**Supplemental Figure 1.**
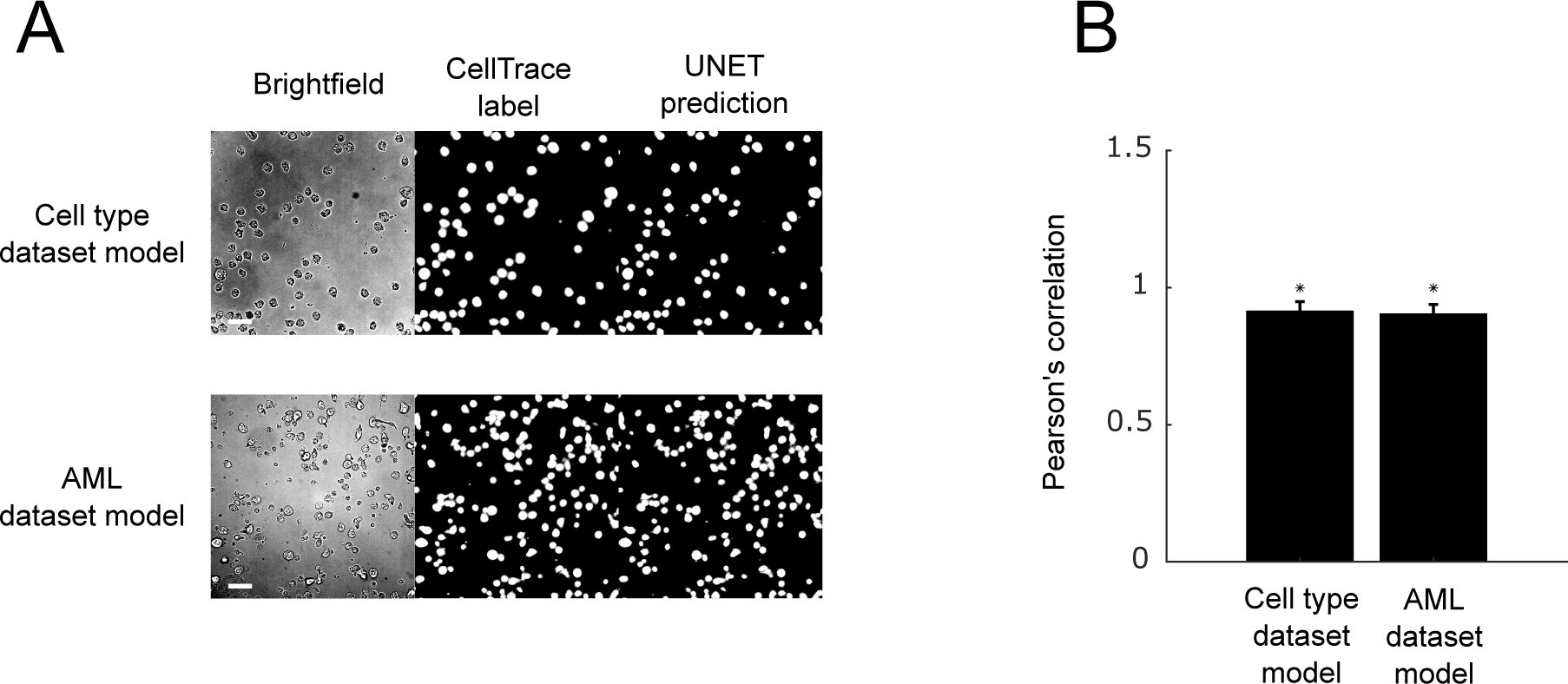
Robust label-free prediction of cell area using the UNET architecture. (A) Sample label-free results from models trained for the cell type dataset and the Acute Myeloid Leukemia dataset. Scale bar represents 20 μm. Brightfield images and respective ground truth fluorescent images are used to compare and calculate Pearson’s correlation values (B) with respect to their predicted synthetic images. * represents the theoretical upper limit of the model’s performance for each dataset. Such a model would perfectly predict the fluorescent level of each cell but not be able to predict fluorescent noises that arise from the instrumentation. (see ^21^ for detailed method) .

**Supplemental Figure 2.**
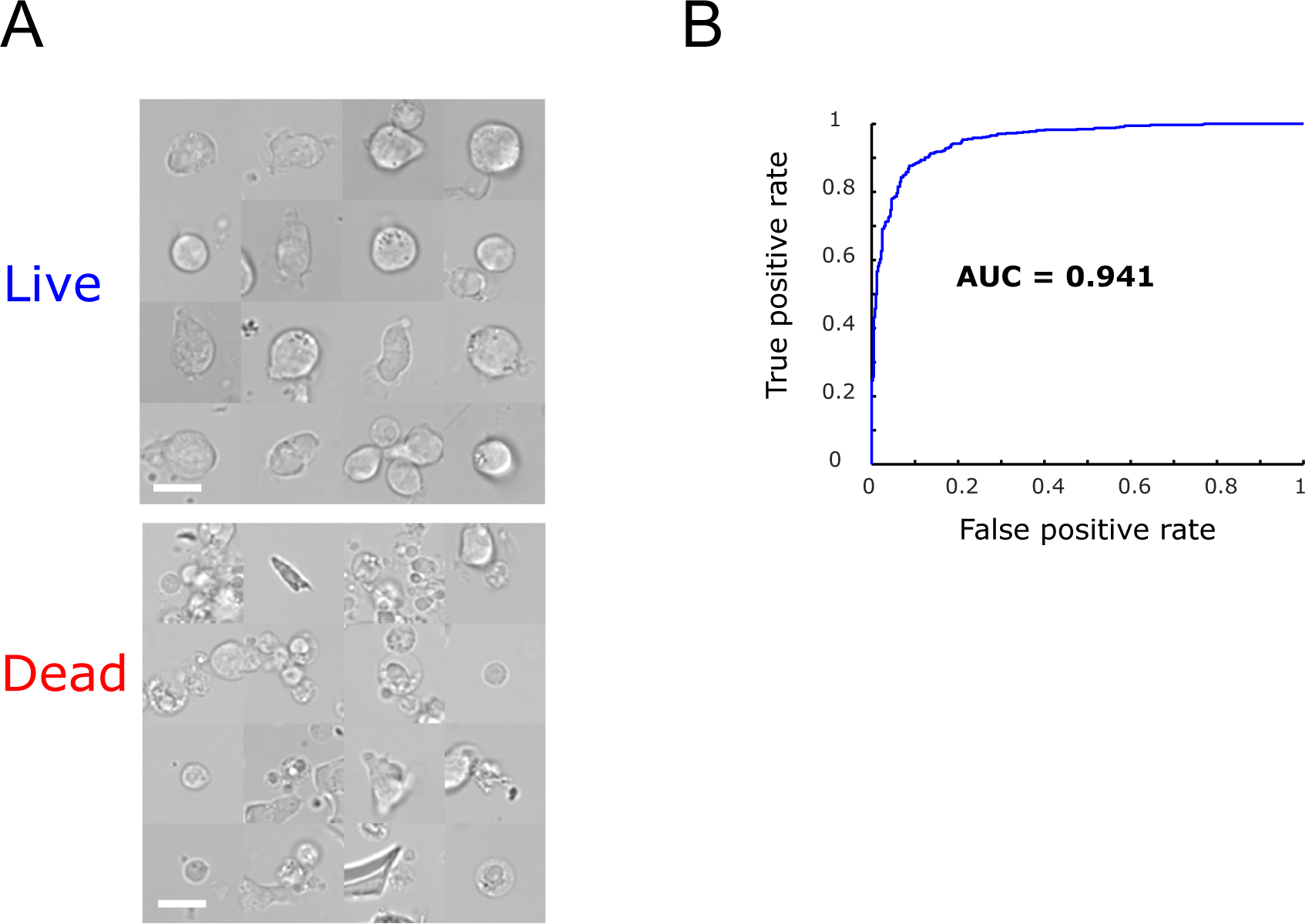
A live dead cell classifier is trained to remove dead cells from the dataset. Brightfield crops of selected cells are labeled as either ‘Live’ or ‘Dead’ using a convolutional classifier that is trained to recognize dead cells using a manually labeled dataset. (A) Representative cell crops classified as ‘Live’ or ‘Dead’. Scale bar represents 10 μm. (B) Receiver operating curve (ROC) measuring the prediction performance of the trained classifier. AUC: Area under the ROC Curve.

**Supplemental Figure 3.**
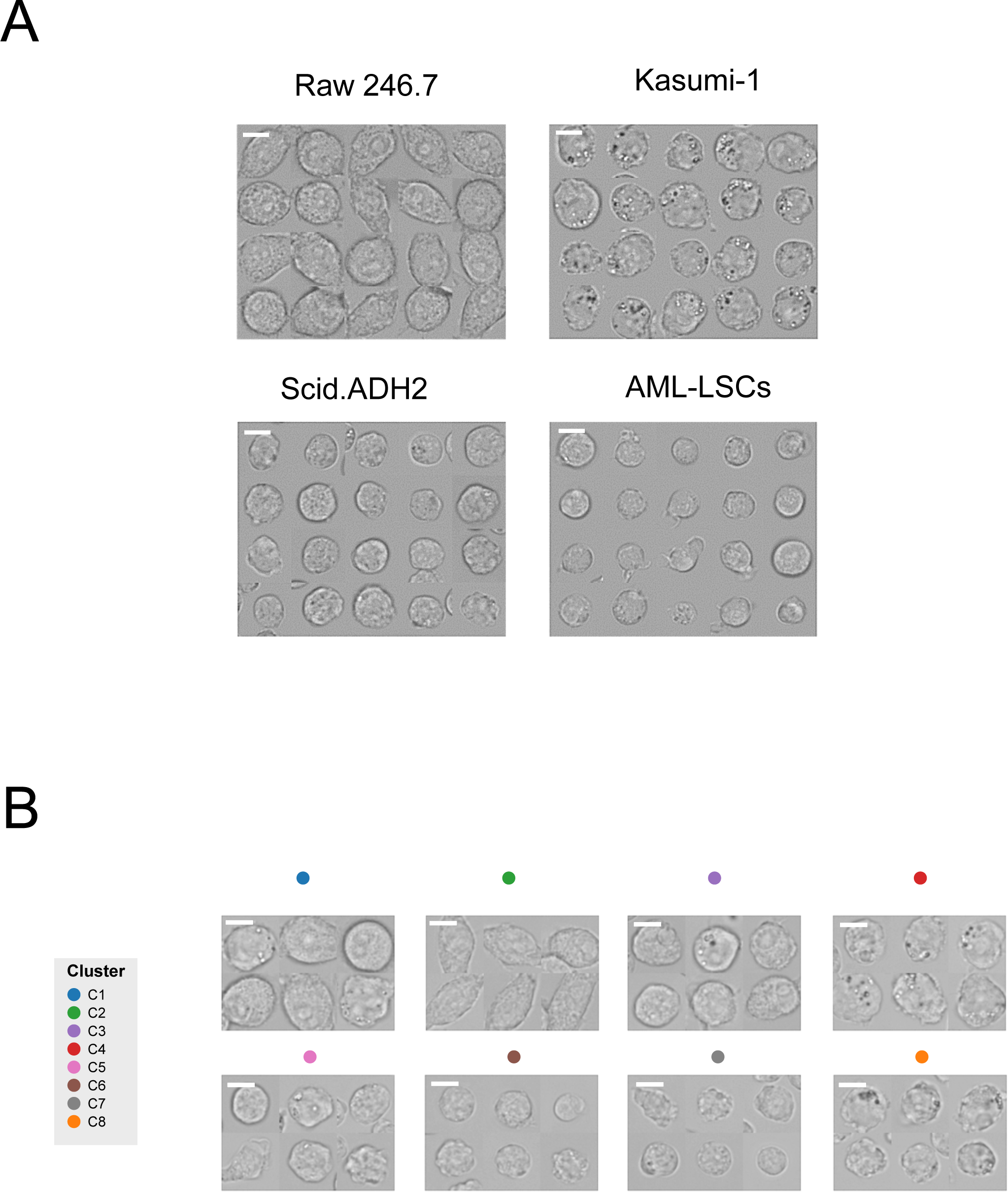
Cell images from blood cell types analyzed by UPSIDE. (A) Representative images from four analyzed blood cell types. (B) Representative images from eight different morphological clusters identified by Louvain clustering of the UPSIDE-generated latent vectors from each cell type. Scale bar represents 5 μm.

**Supplemental Figure 4.**
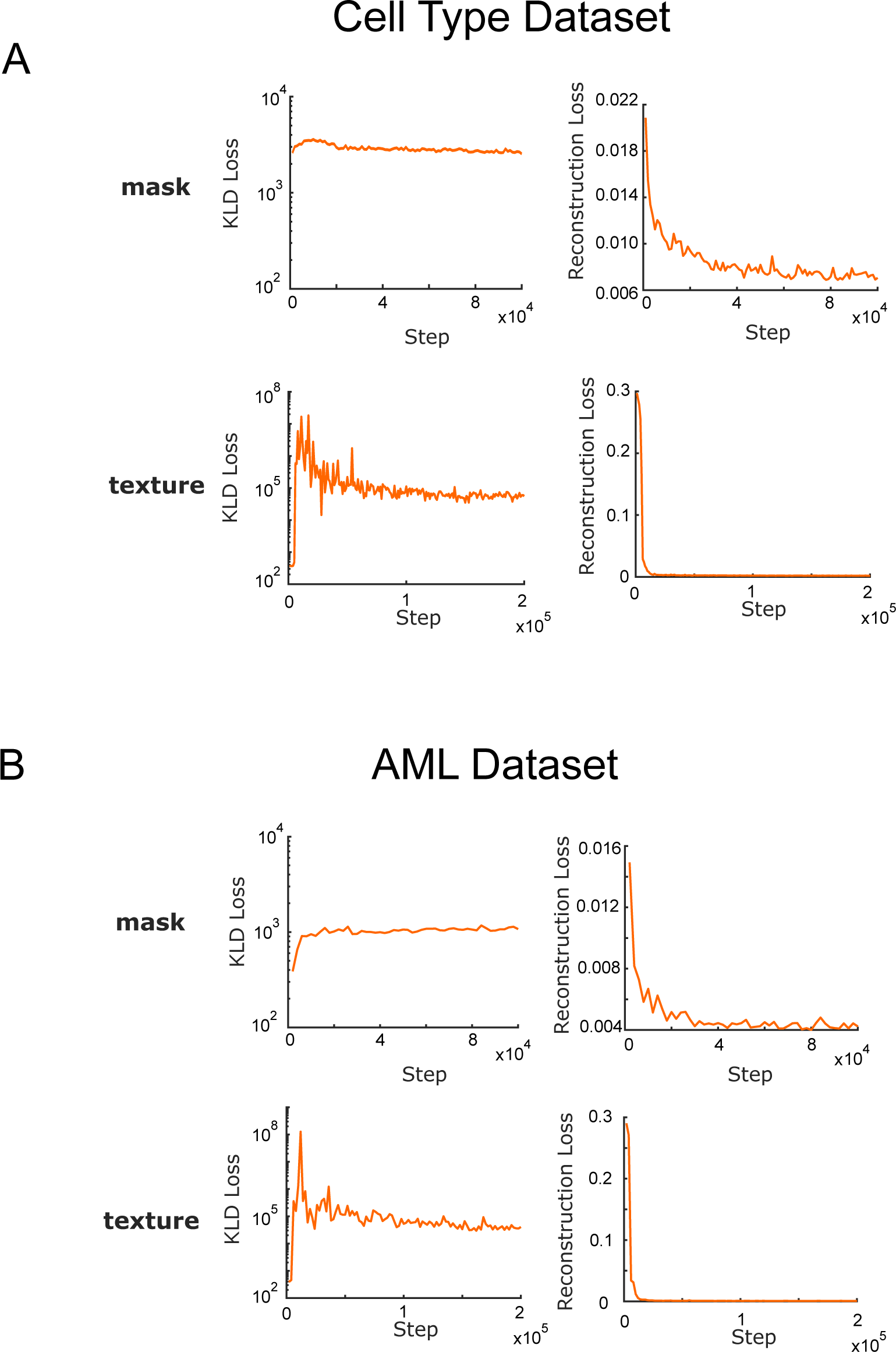
Concurrent training of shape and texture variational autoencoders for the cell type dataset. Reconstruction and Kubeck-Leibler Divergence (KLD) losses of the models for the Cell Types Dataset (A) and the Acute Myeloid Leukemia Differentiation Dataset (B).

**Supplemental Figure 5.**
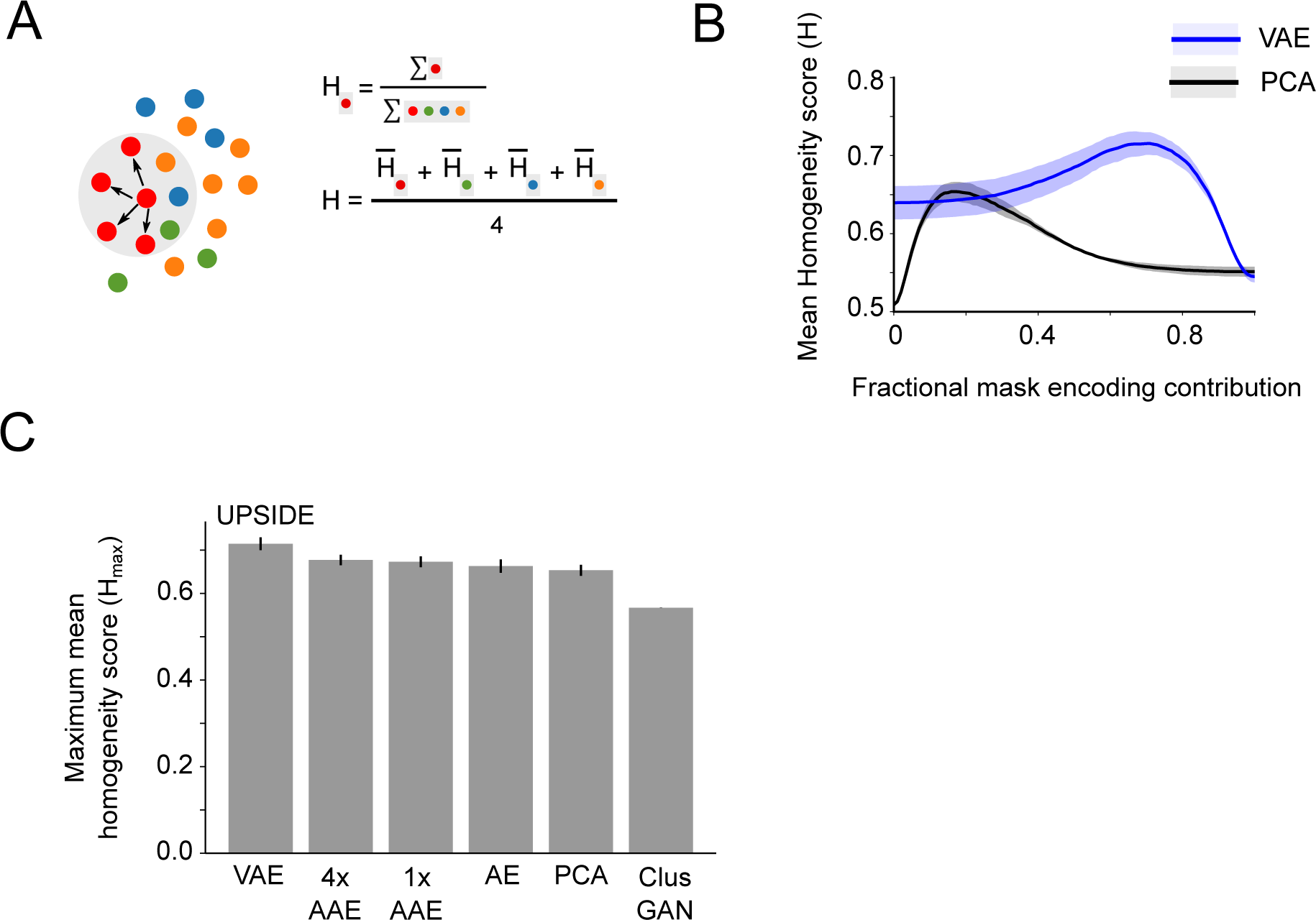
Cell type homogeneity scores were obtained using different data encoding methods. (A) Homogeneity score is calculated to measure how well different cell types are separated in latent space (B) Mean nearest neighbor score (H) across 4 cell types obtained with different relative mask weight contribution for encodings generated by either VAE vs PCA method. (C) Maximum nearest neighbor scores (H_max_) between VAE, PCA, and other alternative deep learning architectures. H_max_ is defined as the highest Mean nearest neighbor score across all weight combinations of mask and texture contributions. VAE: Variational Autoencoder, 4x AAE: Adversarial Autoencoder with latent dimension trained to fit a 4 mixed gaussian distribution, 1x AAE: Adversarial Autoencoder with latent dimension trained to fit a normal distribution, Clus GAN: Cluster Generative Adversarial Autoencoder with the one hot encoding component module removed, PCA: Principal Component Analysis.

**Supplemental Figure 6.**
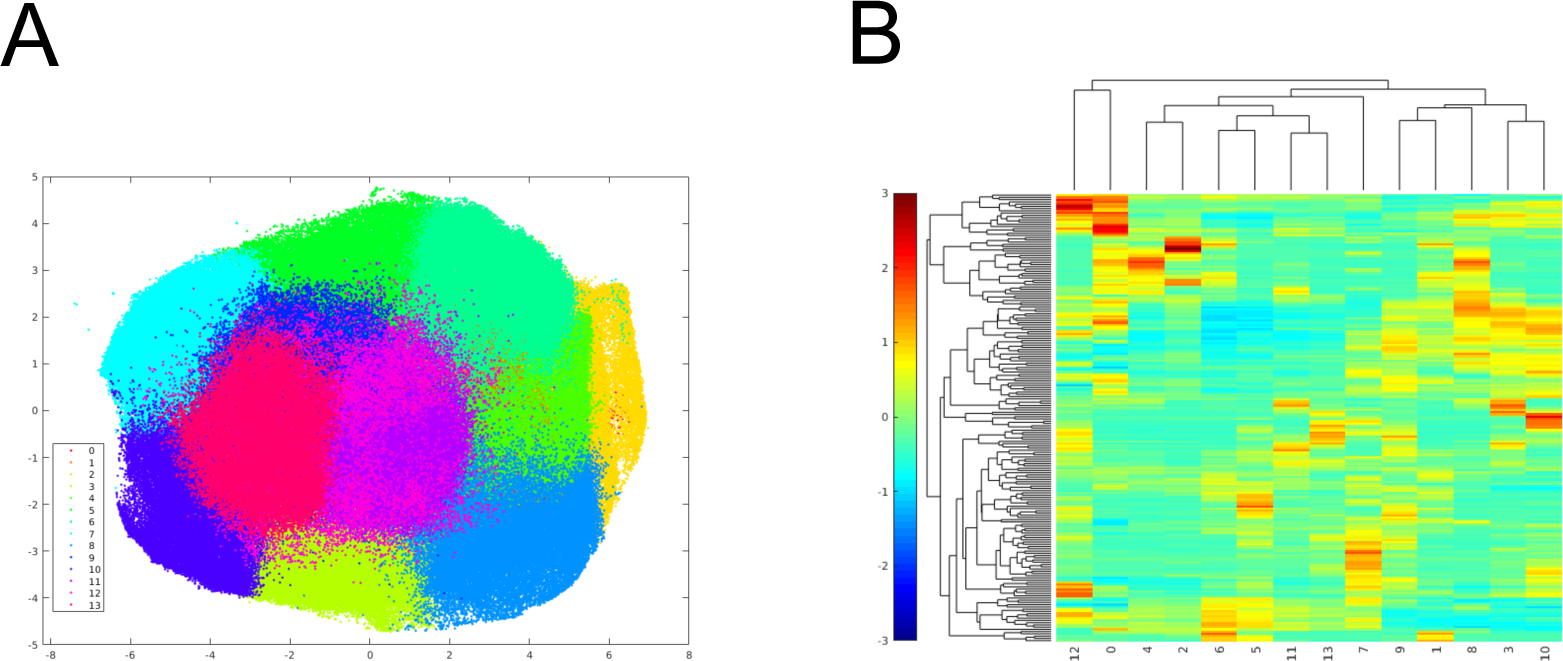
Clustering of AML cell morphologies in the latent space with the Louvain method. (A) 2D UMAP projection of learned mask and texture encodings from combined AML datasets. Each cell was colored based on the raw Louvain clustering result over all datasets. (B) Clustergram of the z-score from morphological groups defined by Louvain methods. Groups with closely related z-score patterns were combined into larger morphological clusters.

**Supplemental Figure 7.**
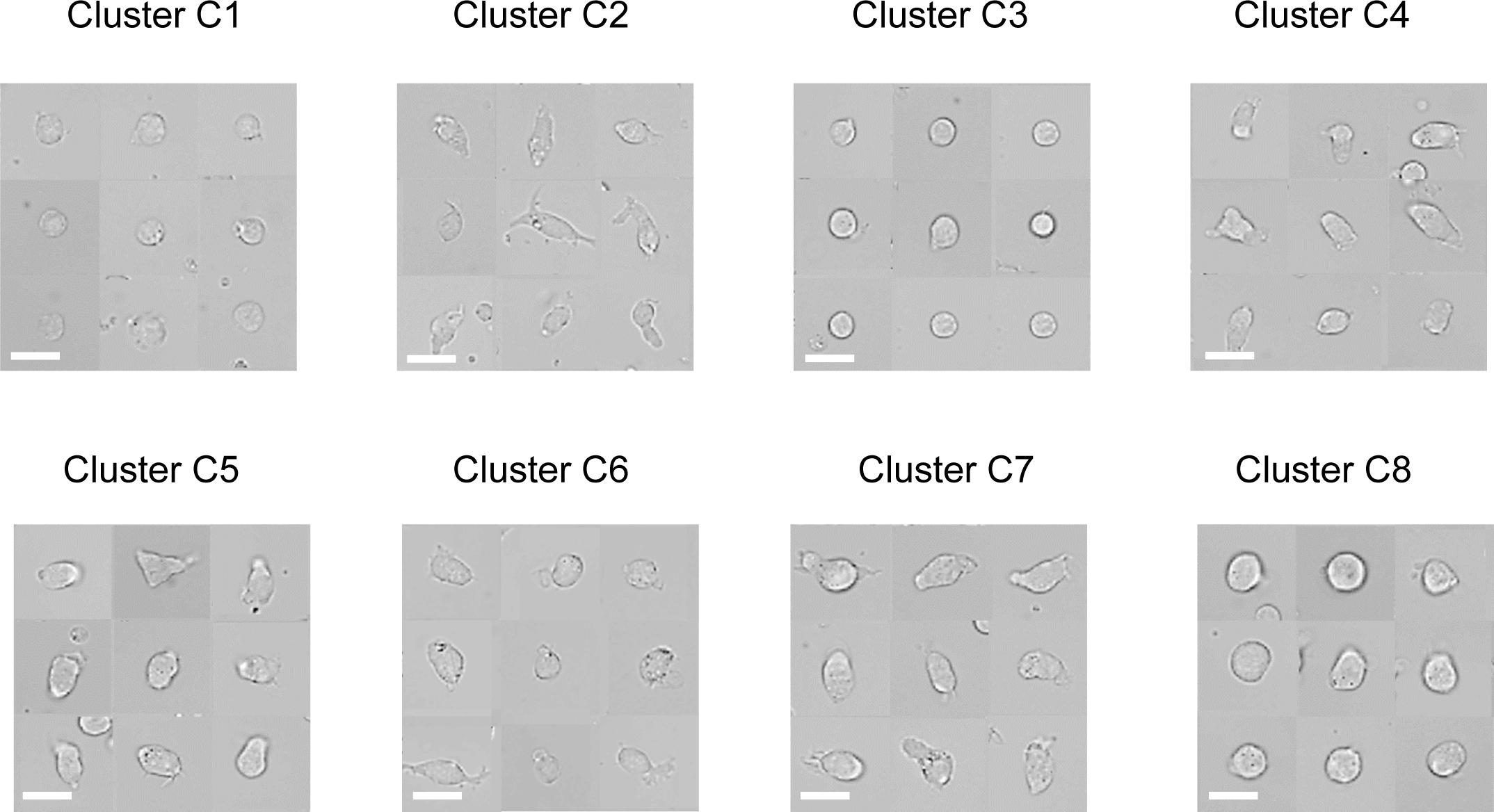
Representative brightfield images of identified cells from each grouped morphological cluster. Scale bar represents 10 μm.

**Supplemental Figure 8.**
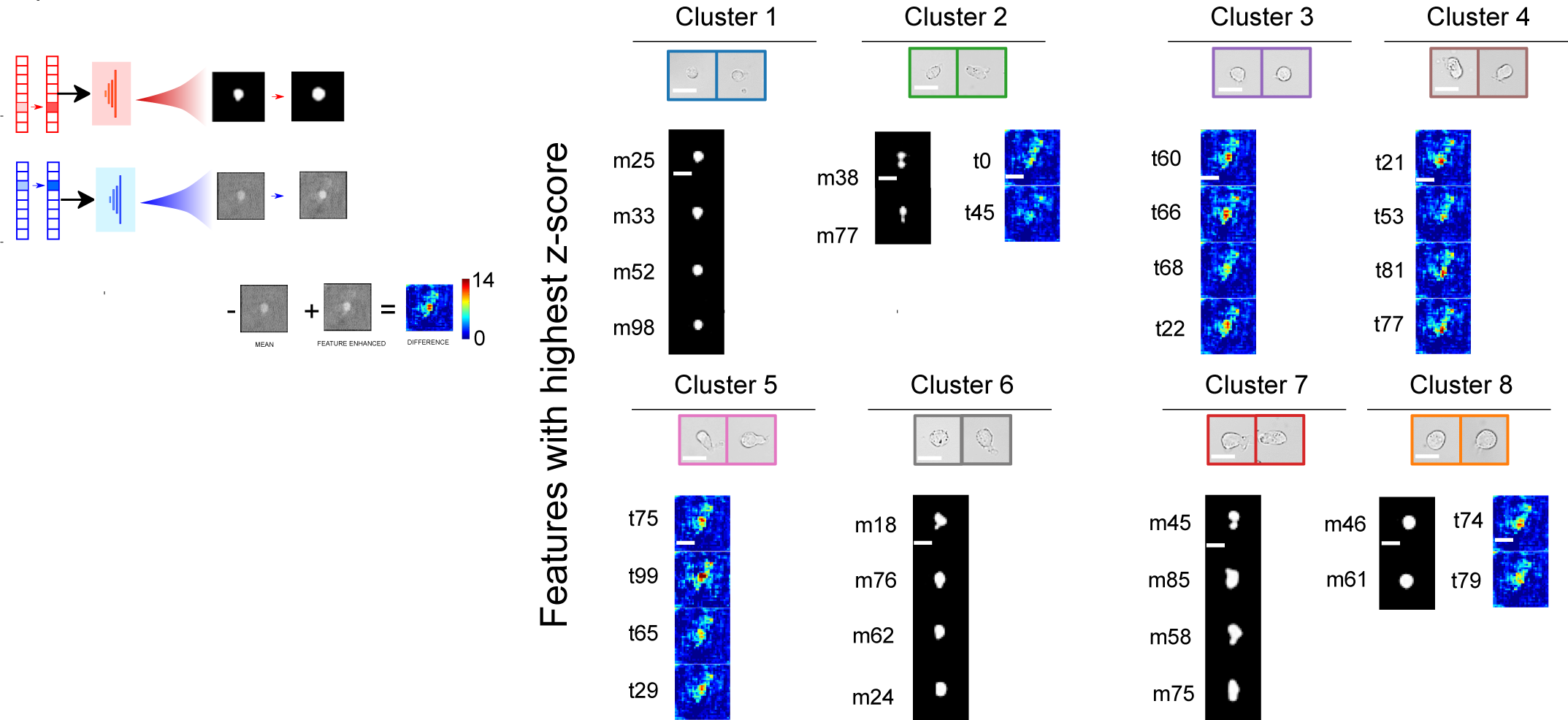
Four most enriched mask and texture features for each morphological cluster are decoded into the image space. Decoded texture images are accompanied by unzoomed pixel difference maps. Scale bar represents 10 μm.

**Supplemental Figure 9.**
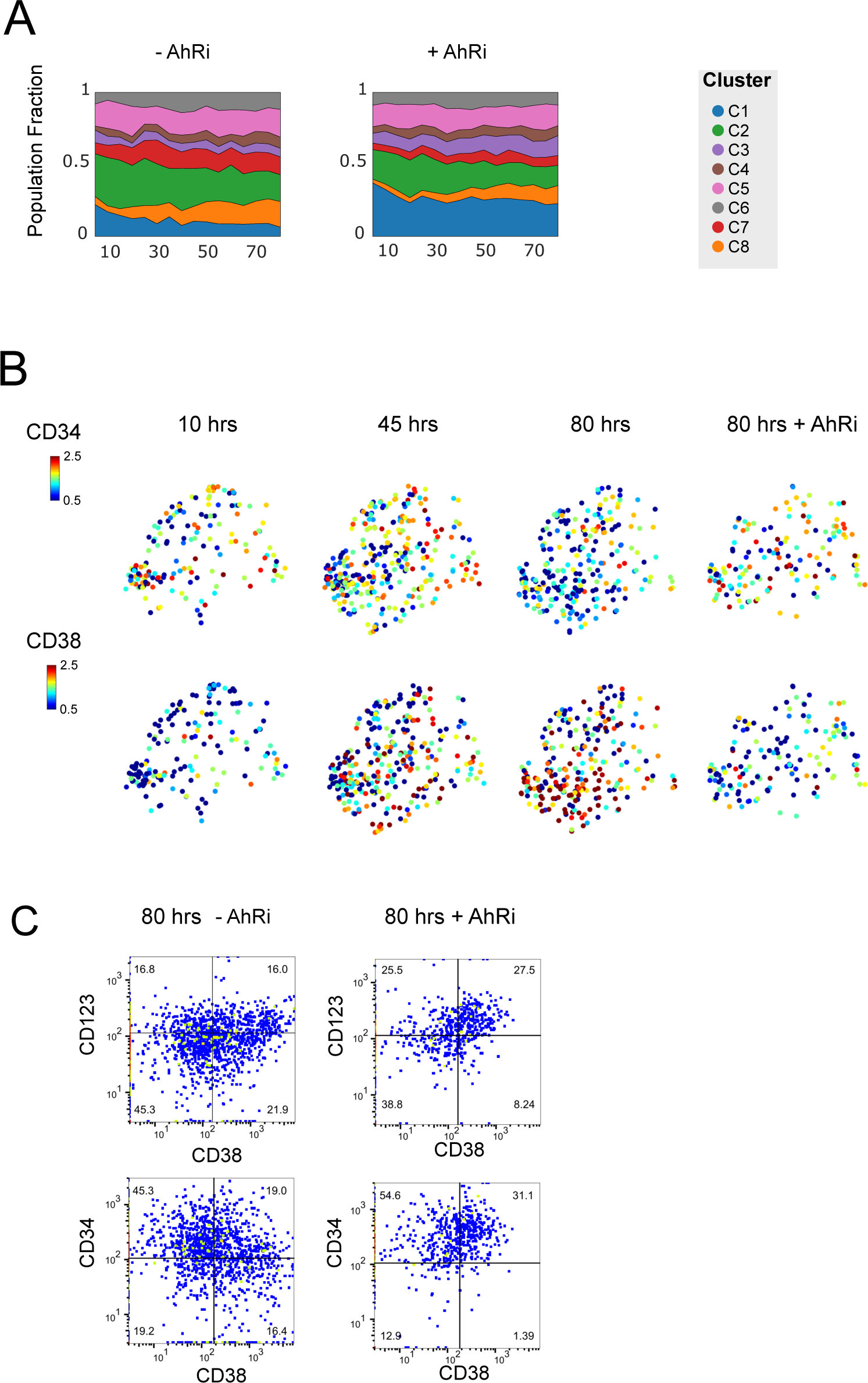
Experimental replication for CD34 and CD38 evolution along with morphological change during LSC differentiation. (A) Dynamics of cells’ CD34 and CD38 expression and their morphological properties during differentiation process in the presence or absence of AhRi. (B) Dynamics of cells’ CD34 and CD38 expression and their morphological properties during differentiation process in the presence or absence of AhRi. (C) Flow cytometry analysis of CD34, CD38, and CD123 expressions of patient LSCs after parallel 80 hrs of non-microscopy culture with or without AhR inhibitors alongside the imaging experiment.

**Supplemental Figure 10.**
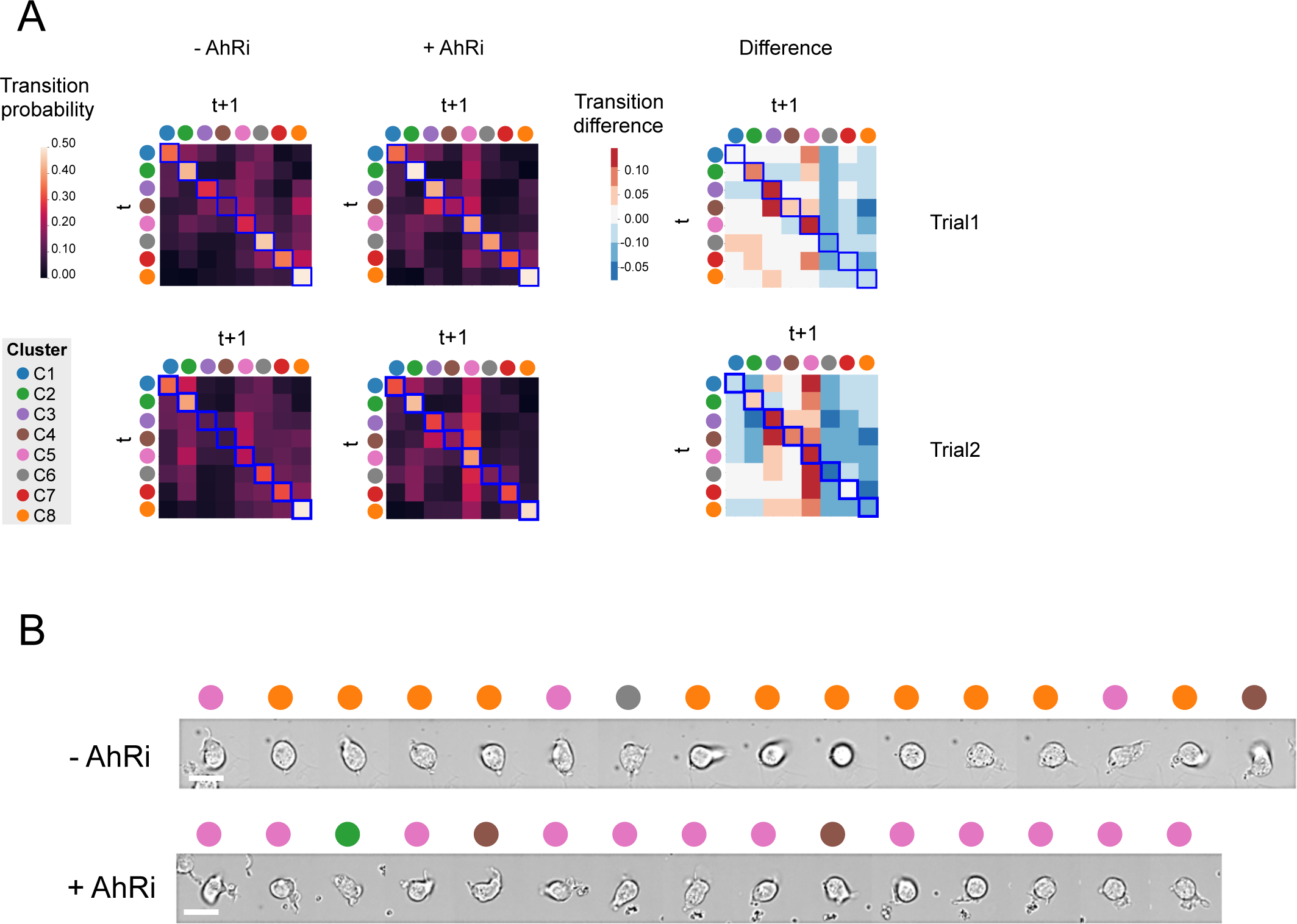
Transition dynamics between morphological clusters for AML in differentiation and stemness-preserving condition. (A) transition probability matrix between identified morphological states with and without AhRi in two replicates (B) Representative cell tracks in cultures with and without AhRi. Scale bar represents 10 μm.

**Supplementary Figure 11.**
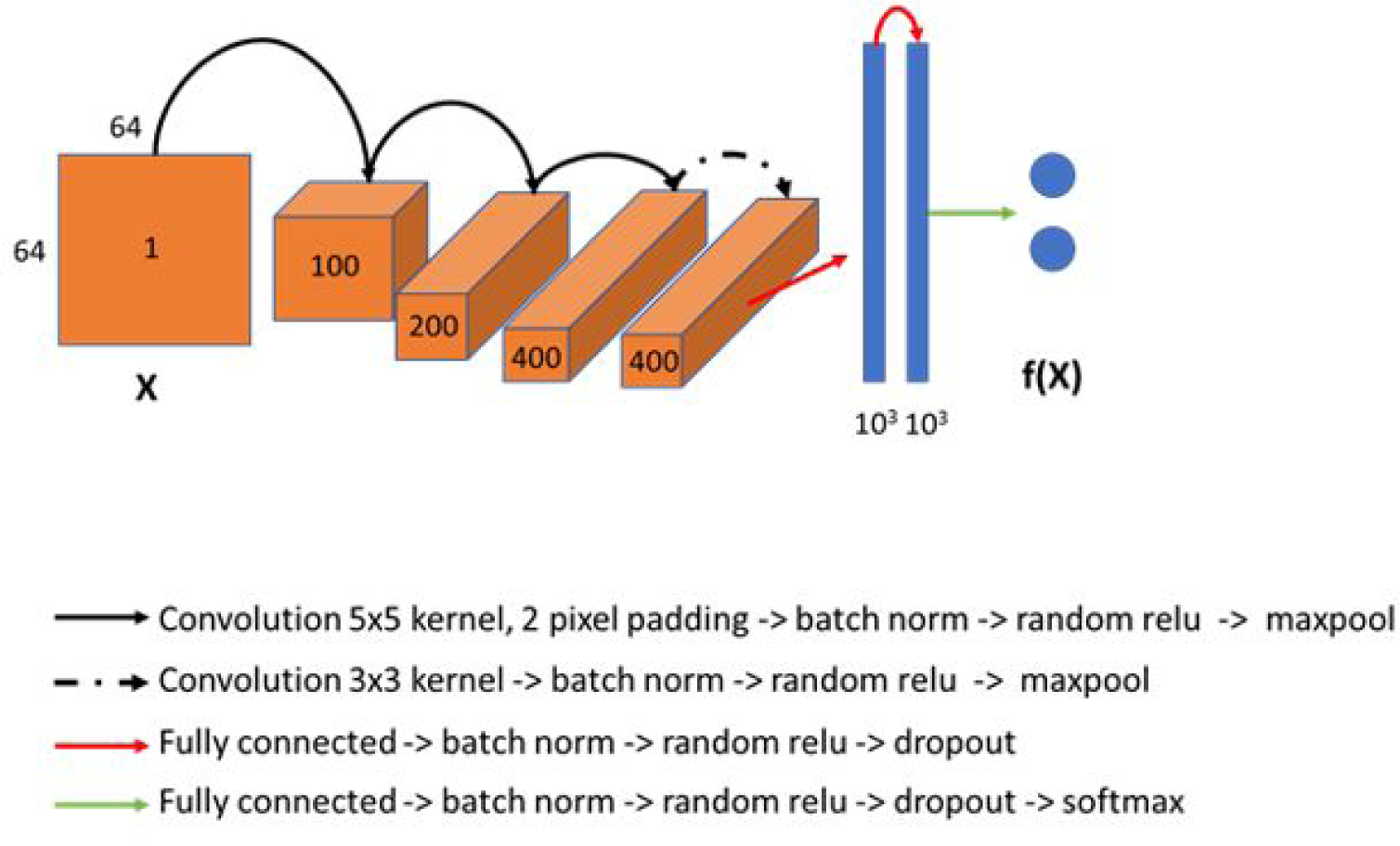
Architecture of convolutional classifier neural network for live cell classification

**Supplementary Figure 12.**
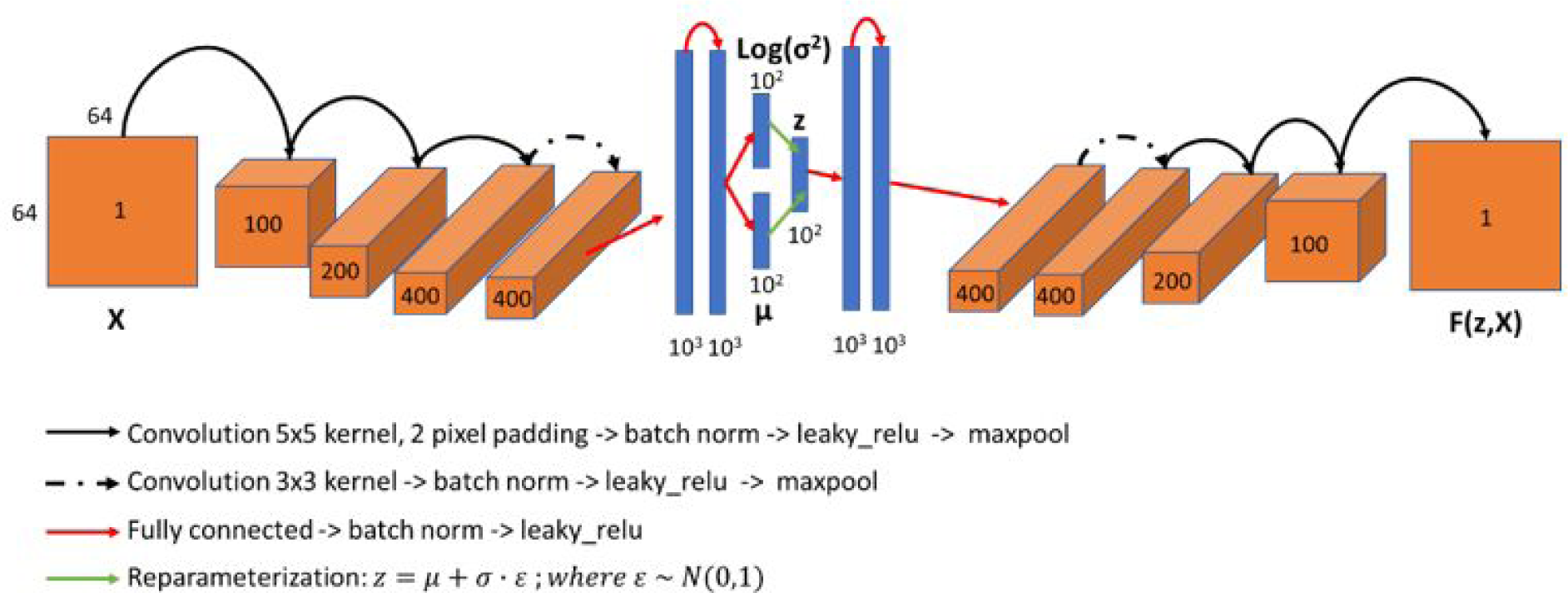
Architecture of convolutional variational autoencoder for cell shape and texture learning.

**Supplementary Figure 13.**
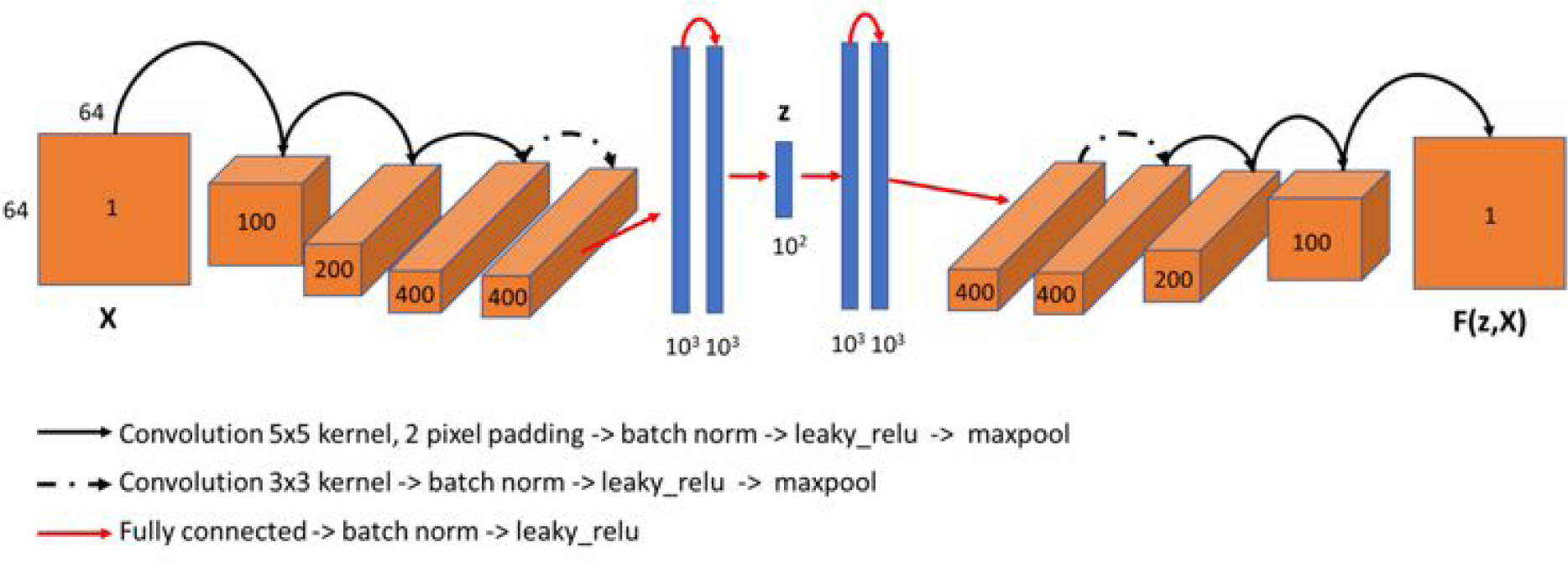
Architecture of convolutional Vanilla Auto Encoder (AE) for cell shape and texture learning.

**Supplementary Figure 14.**
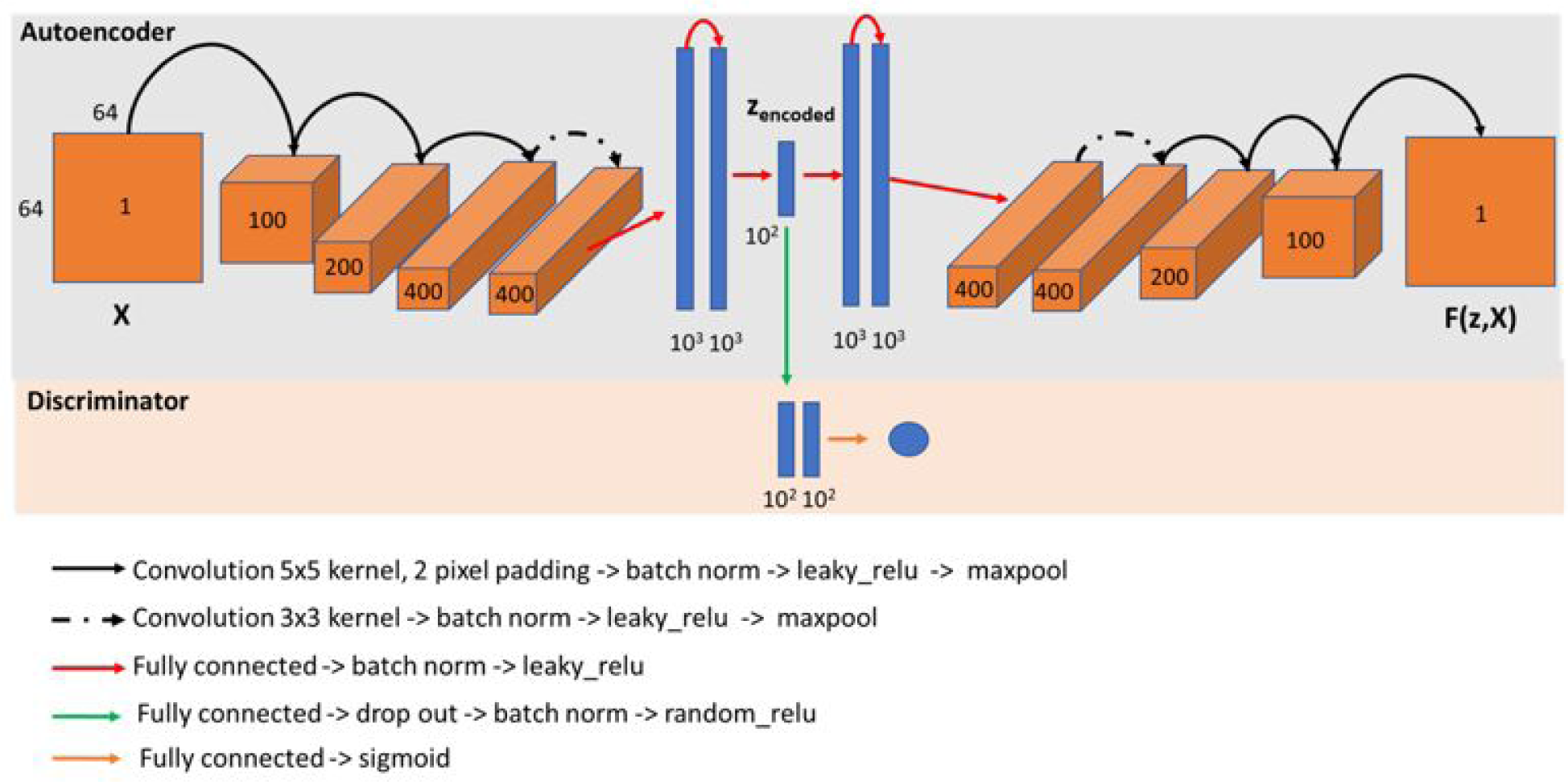
Architecture of convolutional Adversarial AutoEncoder (AAE) for cell shape and texture learning.

**Supplementary Figure 15.**
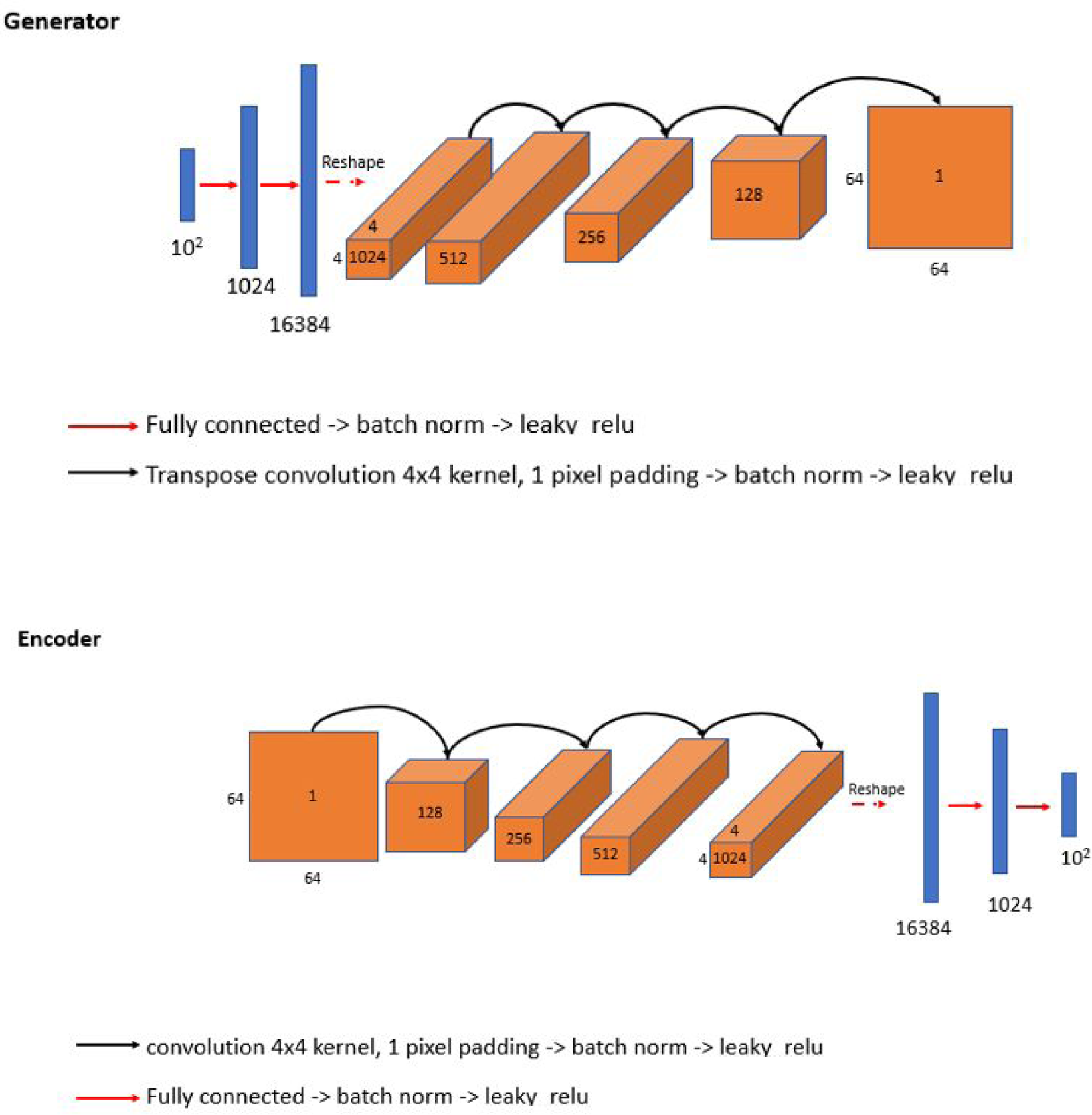
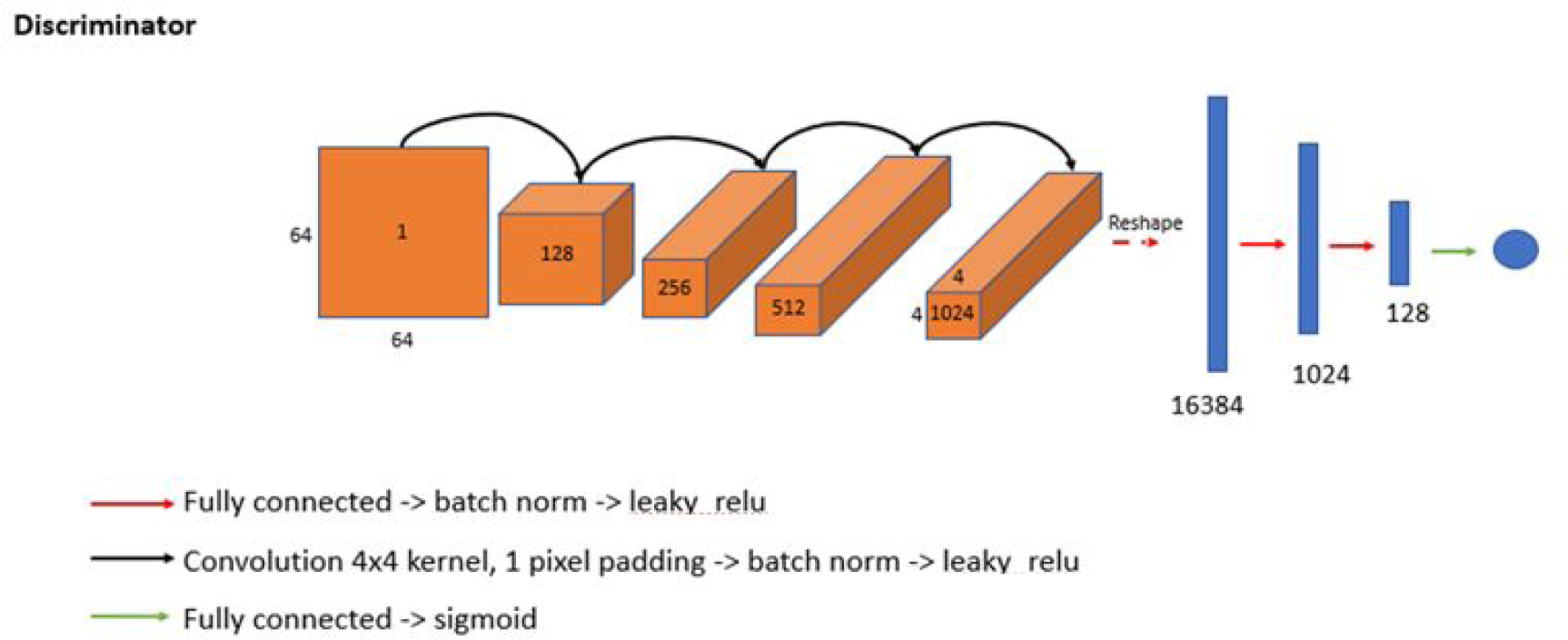
Architectures of the generator, encoder, and discriminator module of clusterGAN for cell shape and texture learning.

**Supplemental Movie 1.**
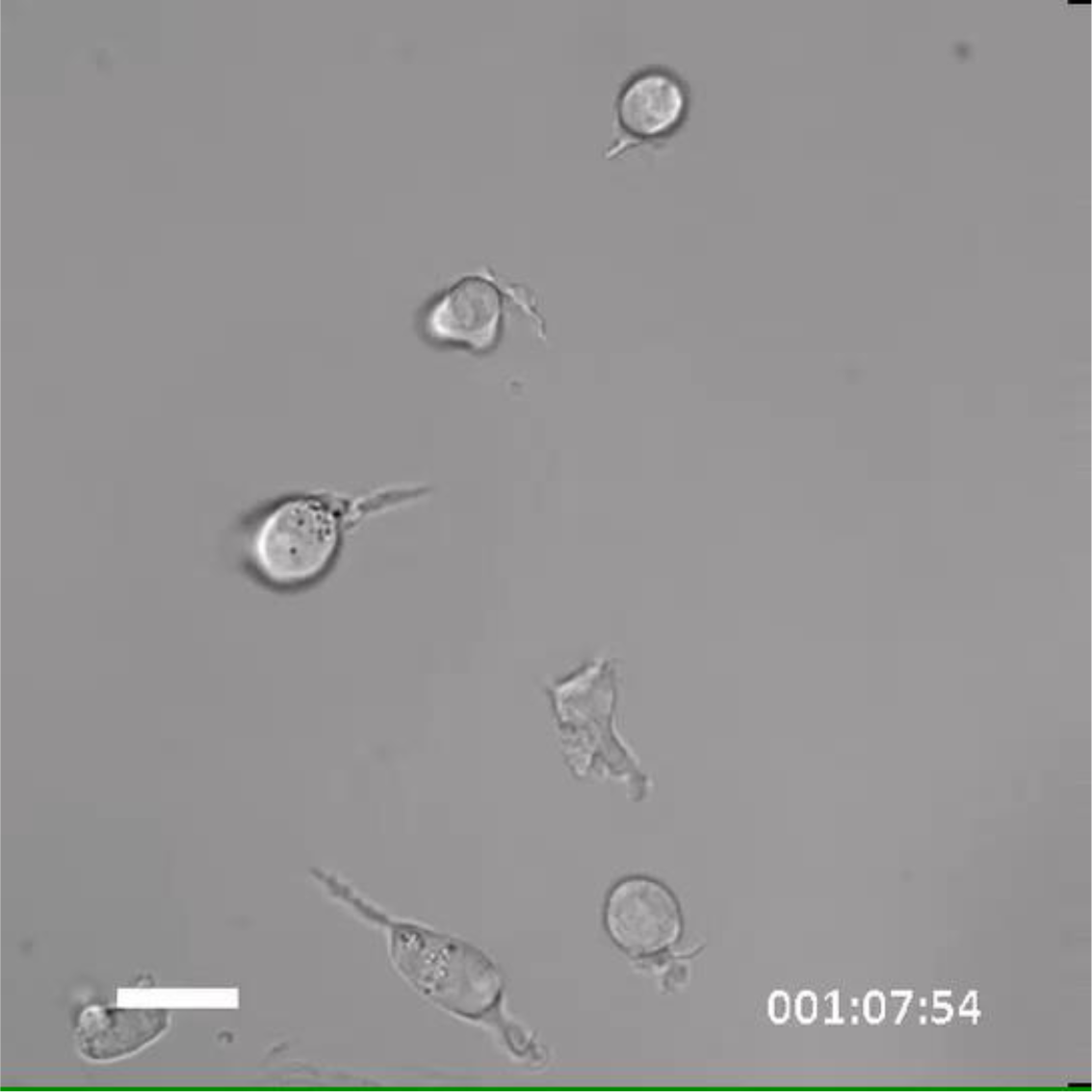
Representative timelapse brightfield movie of cultured AML LSC

